# Ion mobility mass spectrometry and molecular dynamics simulations unravel the conformational stability and coordination dynamics of human metallothionein-2 species

**DOI:** 10.1101/2022.12.01.518729

**Authors:** Manuel David Peris-Díaz, Alexey Barkhanskiy, Ellen Liggett, Perdita Barran, Artur Krężel

## Abstract

Ion mobility-mass spectrometry (IM-MS) unraveled different conformational stability in Zn_4-7_-metallothionein-2. We introduced a new molecular dynamics simulations approach that permitted the exploration of all conformational space confirming the experimental data, and revealed that not only the Zn–S bonds but also α–β domain interaction modulates protein unfolding.

---

Mammalian metallothioneins (MTs) constitute small (~6-7 kDa) cysteine-rich proteins with a primarily biological role in Zn^2+^ and Cu^+^ metabolism.^1–3^ At least a dozen MTs isoforms (MT1-MT4) and multiple subisoforms have been found, which differ in metal-binding properties, tissue, and cellular localization.^4–5^ MTs have proved to be highly challenging objects to study using traditional biophysical techniques due to the lack of secondary structures, aromatic amino acids, and the spectroscopic silence of Zn^2+^.^4^ To date, only one X-ray structure has been solved for rat hepatic mixed Cd_5_Zn_2_MT2 species^6^ The protein adopts a dumbbell shape with two metal-thiolate clusters named α- and β-domains containing a Cd_4_Cys_11_ and Cd_1_Zn_2_Cys_9_ cluster, respectively. Despite Cd^2+^ and Zn^2+^ capabilities to form an M_7_MT2 (where M represents a metal ion), there is a divergence of behavior in the thermodynamic and kinetic properties between these two divalent metal ions, which is not without biological consequences.^7^ The seven Cd^2+^ ions bind cooperatively in Cd_7_MT in a domain fashion with three to five orders of magnitude tighter than Zn^2+^.^2^ In contrast to cadmium, zinc MT2 presents three classes of affinities towards their seven Zn^2+^ ions. Four Zn^2+^ are bound with *K*_d_ ~10^−12^ M, another two with *K*_d_ ~10^−10^-10^−11^ M, while the seventh is weakly bound with *K*_d_ of ~ 10^−8^ M.^8^ As a consequence of this fact, MT2 exists as multiple Zn_4-7_MT2 species under cellular conditions where free Zn^2+^ concentration varies from 10^−11^ to 10^−9^ M.^9–10^ The characterization of isolated MT fractions from several tissues and cell lines supported the role of Zn_4-7_MT2 species as a zinc buffering system.^10–12^ The heterogeneity of these species impedes their study by high-resolution structural techniques like cryo-EM, X-ray crystallography, or NMR. Ion mobility-mass spectrometry (IM-MS) has proven well-suited to interrogate heterogeneous protein systems and characterize their conformation and dynamics. ^13–15^ However, the resolution of the IM device may not be enough to separate closely related protein conformation. In some cases, gas-phase activation of protein ions via collisional activation also referred to Collision Induced Unfolding (CIU)^16^, can be used to probe subtle structural differences between similar conformations and study protein ion stability and dynamics. Recently, high-resolution cyclic IM-MS based on traveling wave technology was introduced, allowing for tandem IM workflows.^17–19^ Despite the structural information that can be derived from IM-MS experiments, it is unable to define protein structure at the atomic level. Recent efforts focused on the integration of molecular dynamics (MD) with IM-MS, assigning gas-phase structures from *in silico* methods.^20–24^ Unfortunately, usually only a few metastable states are explored since biological processes such as protein unfolding or conformational changes are on time scales far beyond those accessible by classical MD simulations. To access other conformational states, as those sampled during CIU, most of the works have used a thermal unfolding approach. While some reports have shown that thermal unfolding can reproduce many general features observed during a CIU experiment^20,24–25^, a recent report has suggested a low reproducibility and lack of conformational sampling.^23^

In order to investigate the conformational properties of Zn_4-7_MT2 species and shed more light on how these species resemble at the microscopic level, IM-MS, CIU, and MD simulations were integrated.

The nESI mass spectra of apoMT2 in the presence of TCEP (termed as “red”) presents a charge state distribution (CSD) spanning only three charge states 3 ≤ z ≤ 5, with apoMT2^5+^_red_ and apoMT2^4+^_red_ the most dominant (**Fig. 1a**). Reconstituted Zn_7_MT2 protein exhibits a CSD shift toward lower charge state with the Zn_7_MT2^4+^_red_ ions predominant (**Fig. 1b**), suggesting in solution-phase conformational changes altering solvent accessible surface area (SASA). The apoMT2^4+^_red_ and Zn_7_MT2^4+^_red_ ions displayed a similar collision cross section (CCS) (**Fig. 1c-d**), hereby not representing the SASA changes. The CCS for apoMT2^5+^_red_ present a broad CCS distribution, and upon Zn^2+^ binding, a single and compact Zn_6_MT2^5+^ and Zn_7_MT2^5+^_red_ conformer is observed (**Fig. S1**). IM-MS revealed that 5+ ions undergo a conformational change capturing the CSD shift observed in the mass-to-charge spectrum (**Fig. 1e-f**). Partially Zn^2+^-loaded MT2 species were obtained via titration in the presence of TCEP, and native IM-MS under different collisional activation (CA) conditions were recorded (**Fig. S2**). Metal-coupled folding effects can be observed upon Zn^2+^ binding to apoMT2^5+^red: prior to CA, Zn_4-7_MT2^5+^_red_ ions populate a CCS ~ 1000 Å^2^ cf. ~ 1300 Å^2^ for apoMT2^5+^_red_ (**Fig. S2**). As the collision energy is increased, the CCS shifts to ~ 1150 Å^2^ in all of the Zn_4-7_MT2 complexes. Similar results were obtained under 50 or 200 mM ammonium acetate (**Fig. S3**). Unpredictable, fitting the native mass spectra to simulated isotopic distributions revealed the partial retention of protons within the Zn^2+^ clusters. Lacking a reducing agent during spraying generates signals shifted to lower *m/z* **(Fig. S2)**. Mass spectra simulations estimated the formation of 7-8 disulfides for all Zn_4-6_MT2^5+^_ox_ and 2 disulfides for Zn_7_MT2^5+^_ox_ (**Table S1**). As a consequence, while some portion of the ions unfolds to ~ 1150 Å^2^, as in the case of reduced complexes, most of the ions are trapped at ~ 1000 Å^2^. To compare the gas-phase stabilities of Zn_4-7_MT2^5+^ ions, the CCS along the CE assayed were fitted to estimate the CIU50 values to indicate the energy required to activate 50% of the ions to its next conformation.^16^ A similar CIU_50_ ~ 90 eV was calculated for Zn_4_MT2^5+^_red_ and Zn_5_MT2^5+^_red_ ions (**Fig. S4**). A gradual increase to CIU_50_ ~ 110 eV was then determined for Zn_6_MT2^5+^_red_ and Zn_7_MT2^5+^_red_. Our previous study provided the location of Zn^2+^ in all Zn_4-7_MT2 species.^26^ In Zn_4_MT2, two Zn^2+^ are bound in each α- and β-domain forming Zn_2_Cys_6_ clusters. The fifth Zn^2+^ binds to the α-domain forming an αZn_3_Cys_9_ cluster. The sixth Zn^2+^ saturates the α-domain forming an αZn_4_Cys_11_ cluster, and the seventh Zn^2+^ forms the βZn_3_Cys_9_ cluster. CIU did not detect structural changes between Zn_2_Cys_6_ and Zn_3_Cys_9_ clusters but determined elevated structural stability upon the formation of the Zn_4_Cys_11_ cluster. Taken together, our results elucidated a plausible structural explanation for why the seventh Zn^2+^ ion binds with a *K*_d_ of ~10^−8^ M to MT2 and provided a link between structural and Zn^2+^ buffering properties.^8^ To get further structural insights into the existing conformational families, we employed multistage IM-MS using a cyclic IM-MS instrument. Upon activation of isolated compact conformer α, we observe an unfolding profile that leads largely to a conformation γ, through β intermediate conformation and a minor extended δ conformation (**Fig. 2a-c**). The great potential of tandem IM is that it allows not only to examine unfolding mechanisms but also to evaluate of the thermodynamic and kinetic stability of unfolded conformations.^18^ Isolation of the conformer γ (**Fig. 2d**) and subsequent CIU activation leads to an unfolding profile with no new features (**Fig. 2e**). We do not evidence a sign of interconversion to a compact conformation (**Fig. 2f**). These results suggest that exists a relatively high transition energy barrier between these states, and propose a plausible irreversible unfolding mechanism that yields to a thermodynamically stable conformation that can be of relevant interest under cellular stress conditions. To understand the experimental results from a microscopic point of view, we then performed gas-phase MD simulations **(Fig. 3a)**. We first focused on examining how the Zn_4-7_MT2 protein complexes are transferred from the solution into the gas-phase (**Fig. S5-S10, 3b**). Proteins were placed in aqueous nanodroplets with an excess of charge to 16+ using Na^+^ as a charge carrier rather than H^+^. As the simulation evolves, the solvent gradually evaporates, and Na^+^ ions are ejected until a charge decrease to 5+. Droplet shrinkage is accompanied by a decrease in the CCS from ~ 1200 to 1000 Å^2^ until the structure collapse to a more compact conformation with CCS ~ 950 Å^2^ without altering the Zn–S bonds **(Fig. S5-S10, 3c)**. In this process, salt bridges do not appear to modulate the transition from solution to gas-phase^27^, but the collapse of the structure is highlighted by an increased number of h-bonds from ~20 to ~40 **(Fig. 3d-e)**. Such results are in excellent agreement with our experimental native IM-MS data (max error < 5 %) and shed light on the gas-phase desolvation and protein structure at the atomic level. We then attempt to simulate the CIU of electrosprayed protein complexes. First, a thermal unfolding protocol was performed on representative structures obtained from gas-phase MD desolvation simulations **(Fig. 3f)**. All of the protein ions have narrower ΔCCS than recorded experimental CIU data **(Fig. 3a)**: apoMT2^5+^_red_ has a ΔCCS of 400 Å^2^ vs. ΔCCS 300 Å^2^ for simulations, and Zn_4-7_MT2^5+^_red_ have a ΔCCS of 200-250 Å^2^ vs. ΔCCS 70-150 Å^2^ for simulated proteins. SA identified multiple conformations for apoMT2^5+^, a compact conformation ~ 1000 Å^2^, and a semi-extended ~ 1200 Å^2^, although the extended one with CCS ~ 1300 Å^2^ was not sampled **(Fig. 3f)**. Similarly, the compact and semi-extended conformers were present in αZn_2_βZn_2_MT2^5+^ and αZn_3_βZn_1_MT2^5+^ with histograms displaying a bimodal distribution, and once again, the extended conformation was not detected. As the Zn^2+^ loading increases, the conformational heterogeneity measured as the CCS (ΔCCS) decreases, indicating that Zn^2+^ promotes protein folding. Consequently, simulations for Zn_5_MT2^5+^ and higher Zn^2+^-loaded states only sampled the compact conformations. Extending the simulation times from 10 to 100 ns did not influence the conformational sampling. Therefore, thermal unfolding failed to overcome the energetic restraints imposed by the Zn–S bonds and was not able to sample extended conformations obtained upon collisional activation. As the radius of gyration (*R*_g_) correlated well with the CCS values, we used an enhanced sampling algorithm named steered MD (SMD) simulations to accelerate the transitions between different states by using *R*_g_ as a collective variable (CV). The force-CCS profiles obtained by SMD simulations clearly distinguished the compact α and semi-extended conformation β but also sampled the extended conformation γ in all cases, and reproduce well their the ΔCCS **(Fig. 3g, S11-S17)**. Comparable CCS distributions were obtained when using the end-to-end N-C terminus distance as a CV **(Fig. S18-S24)**. We observed that protein unfolding proceeds via destabilizing the interdomain α–β interactions (**Fig. 3h**). To date, no single report has characterized the structure and protein conformations for physiologically relevant zinc MT2 species. Here IM-MS aided by MD simulations present a comprehensive structural characterization of these protein complexes. Collectively, our study provides a plausible link between structural and Zn^2+^ buffering properties and sheds light on the mode of functioning of these small yet critical cellular proteins. In addition, as thermal unfolding has been shown not only here but also in other studies that may lack conformational sampling, we report an alternative MD framework to simulate CIU experiments.

**Fig. 1.**
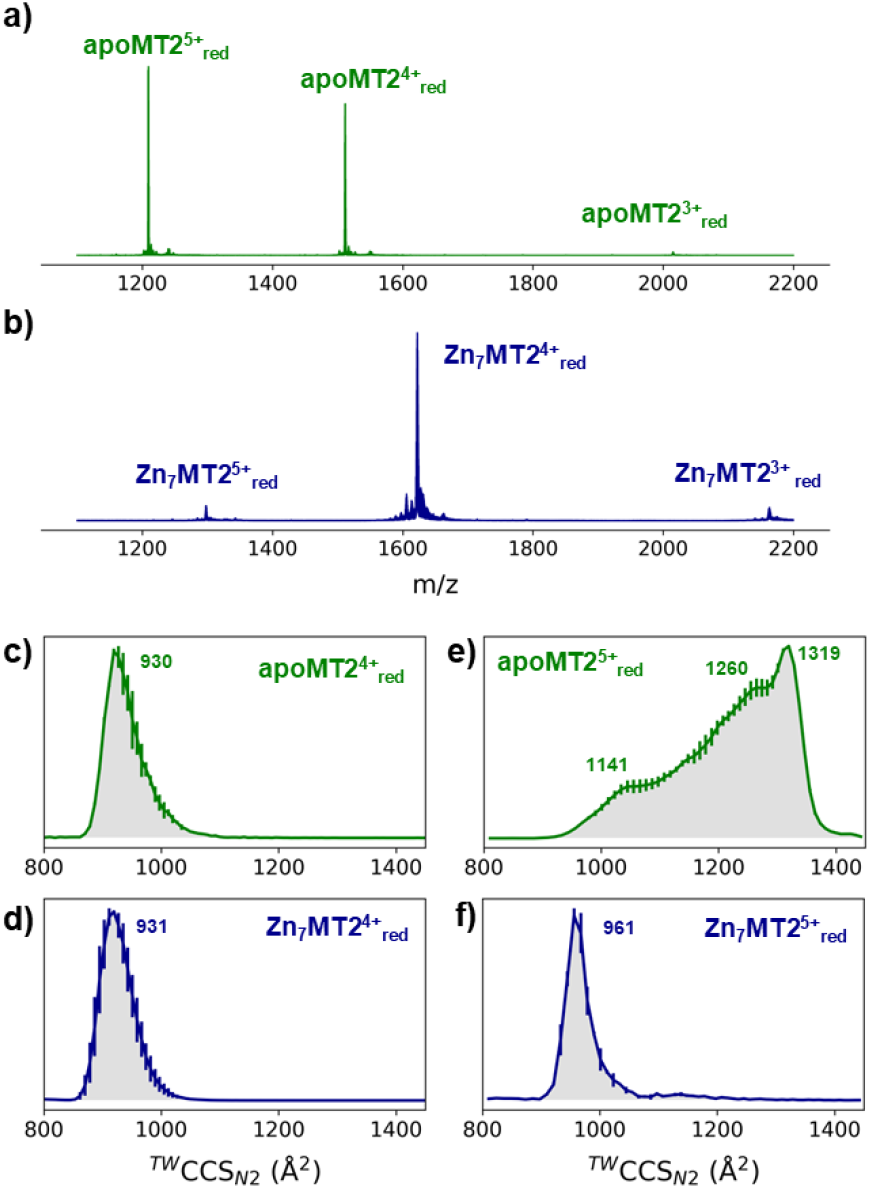
Native mass spectra of apoMT2 and Zn_7_MT2 (a-b) and travelling wave (TW) ion mobility (IM)-derived collision cross sections (CCS) (c-f) of quadrupole-selected apoMT2 and Zn_7_MT2 5+ and 4+ ions. The proteins (10 μM) were sprayed in 50 mM ammonium acetate (pH 6.8) in the presence and absence of 1 mM neutralized TCEP (pH 7.4). Red refers to the reduced state of the ions, as TCEP was on-line employed during the measurement. The CCS values were calculated from three replicates, and the error bars plot along the CCS axis.

**Fig. 2.**
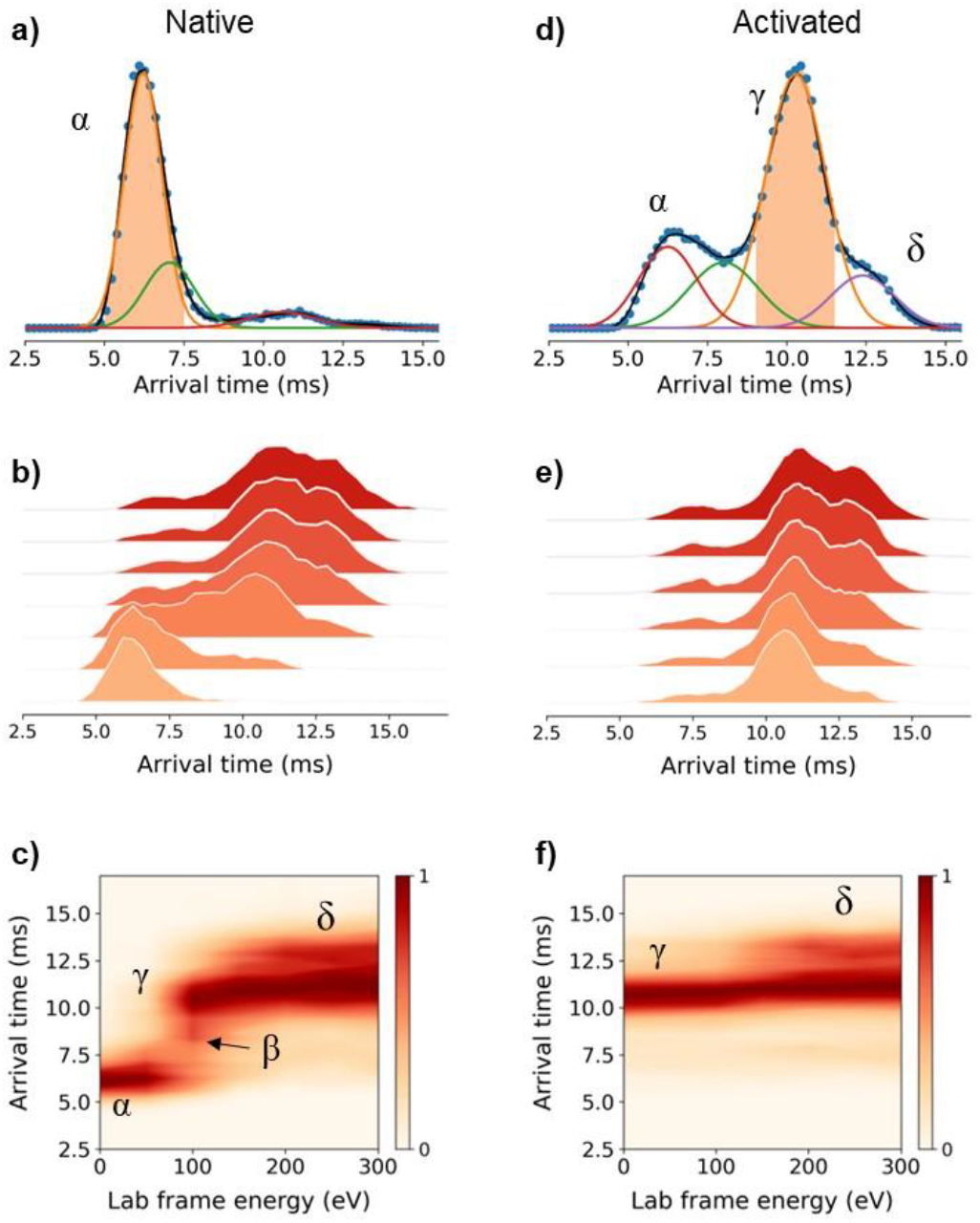
Multistage cyclic IM-MS experiments. Arrival time distribution (ATD) recorded for the mass-selected Zn_7_MT2^5+^_red_ ions (1298 *m/z*) under non-activating conditions (a) and activated on injection to the trap prior to IM selection (d). IMS-CA-IMS and CA-IMS-CA-IMS approach in which the conformer α (b) and conformer γ (e) were isolated and reinjected from the pre-store into the array at increasing activation energies, respectively. Unfolding profiles for the IMS-CA-IMS (c) and CA-IMS-CA-IMS approach (f).

**Fig. 3.**
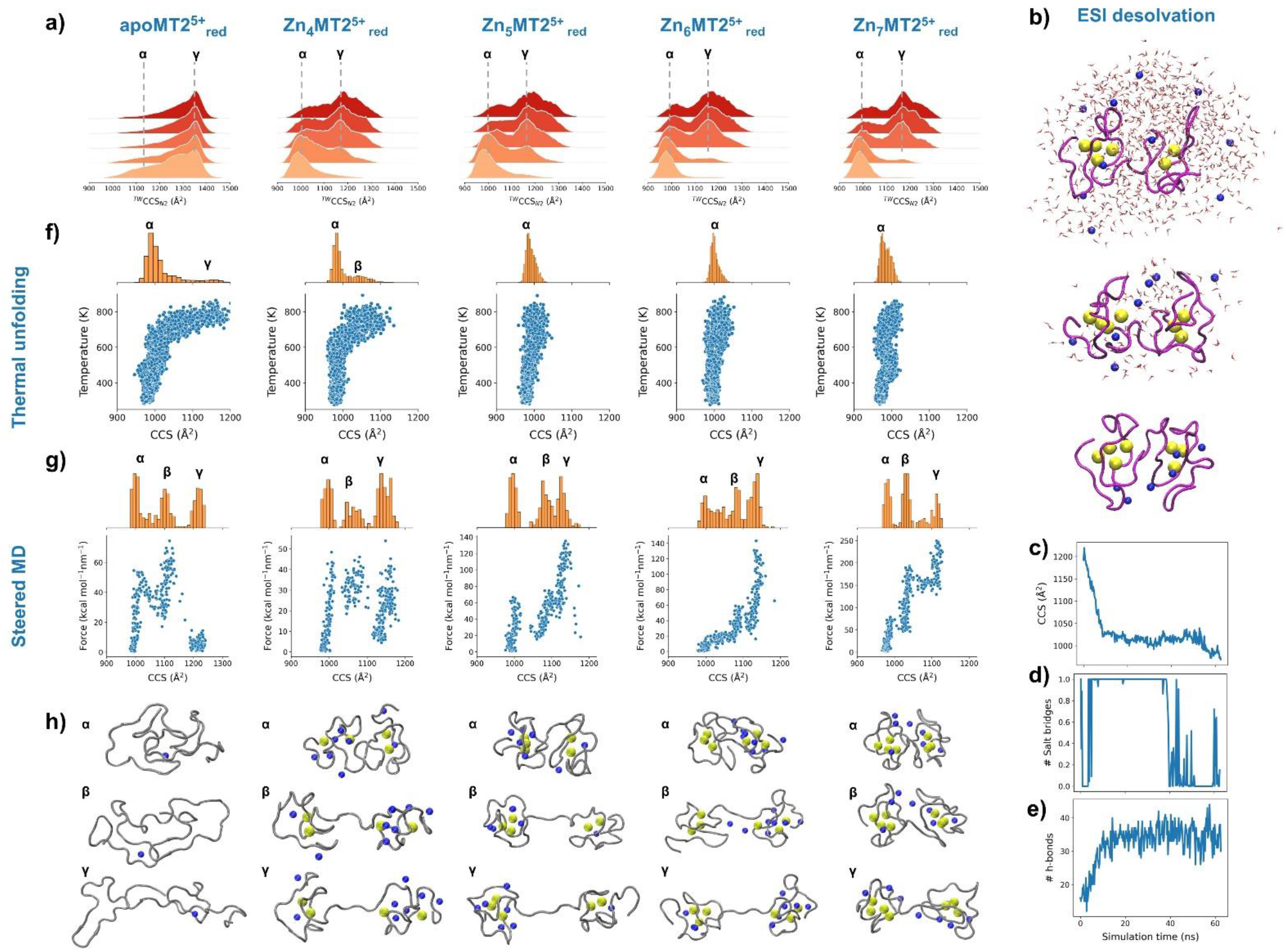
Collision cross sections (CCS) profiles for the quadrupole-selected apoMT2^5+^_red_ and Zn_4-7_MT2^5+^_red_ species under different collisional activation energies (a). Snapshots of the desolvation process at different simulation times for an aqueous nanodroplet containing Zn_7_MT2 and Na^+^ as charge carrier (b) and analysis of CCS (c), salt-bridges (d) and h-bonds (e) as a function of desolvation time. CCS-Temperature plots obtained from thermal unfolding simulations (f). CCS-Force plots derived from SMD simulations (g) and representative snapshots for the conformations from each protein species labeled as in the CCS histograms (h). Yellow and blue spheres represent Zn^2+^ and Na^+^, respectively, and the solvent molecules’ oxygen atoms are shown in red. The protein backbone is shown in magenta or grey for simulation of the ESI or CIU process, respectively.

This research was supported by the National Science Centre of Poland (NCN) under the Opus grant no. 2018/31/B/NZ1/00567 (to A.K.), Preludium no. 2018/31/N/ST4/01909 and Etiuda no. 2020/36/T/ST4/00404 (to. M.D.P.D). We acknowledge the support of EPSRC through the strategic equipment award EP/T019328/1, the European Research Council for funding the MS SPIDOC H2020-FETOPEN-1-2016-2017-801406 and Waters Corporation for their continued support of mass spectrometry research within the Michael Barber Centre for Collaborative Mass Spectrometry.

