## Supporting Information for "Ion mobility mass spectrometry and molecular dynamics simulations unravel the conformational stability and coordination dynamics of human metallothionein-2 species"

*Electronic Supporting information*

**TABLE OF CONTENTS**

Materials ..................... S2
Expression and purification of metallothioneins..................... S2
Mass spectrometry. ..................... S3
Computational studies..................... S6
Figure S1..................... S11
Figure S2..................... S12
Figure S3..................... S12
Figure S4..................... S13
Figure S5..................... S14
Figure S6..................... S16
Figure S7..................... S17
Figure S8..................... S18
Figure S9..................... S19
Figure S10..................... S20
Figure S11..................... S21
Figure S12..................... S22
Figure S13..................... S23
Figure S14..................... S24
Figure S15..................... S20
Figure S16..................... S21
Figure S17..................... S22
Figure S18..................... S23
Figure S19..................... S24
Figure S20..................... S30
Figure S21..................... S31
Figure S22..................... S32
Figure S23..................... S33

Figure S24 ........ S34
Table S1 ........ S35
Table S2 ........ S36
REFERENCES ........ S37

### EXPERIMENTAL SECTION

**Materials.** The following reagents: $ZnSO_4 \cdot 7H_2O$, 4-(2-pyridylazo)resorcinol (PAR), $(NH_4)_2CO_3$, tris(hydroxymethyl)aminomethane (Tris base) and 4-(2-hydroxyethyl)-1 piperazineethanesulfonic acid (HEPES), mass spectrometry grade methanol, tris(2carboxyethyl)phosphine hydrochloride (TCEP), ammonium acetate (AmAc) ethylenediamine-tetraacetic acid (EDTA), and mass spectrometry grade acetonitrile (ACN) were purchased from Sigma-Aldrich. Resin Chelex 100 was acquired from Bio-Rad and 98% hydrochloric acid (HCl) was purchased from VWR Chemicals. Dithiothreitol (DTT) was purchased from Iris Biotech GmbH. Tryptone, LB broth, yeast extract, isopropyl-β-D-1-thiogalactopyranoside (IPTG), and SDS were from Lab Empire, NaCl, NaOH, glycerol, $KH_2PO_4 \cdot H_2O$, $K_2HPO_4$ from POCH (Gliwice Poland), pTYB21 vector and chitin resin were from New England BioLabs, and 5,5′-dithiobis-(2-nitrobenzoic acid) (DTNB) from TCI Europe N.V. was purchased from Sigma-Aldrich.

**Expression and purification of metallothioneins**. Expression vector (Addgene plasmid ID 105693) were transformed into BL21(DE3) *E. coli* cells and growth in culture medium (1.1% tryptone, 2.2% yeast extract, 0.45% glycerol, 1.3% $K_2HPO_4$, 0.38% $KH_2PO_4$) at 37°C until ~0.5 $OD_{600}$. Protein was induced by adding 0.1 mM IPTG to cells and overnight incubation at 20°C with vigorous shaking. The next steps were conducted at 4°C. Cells were centrifuged (4,000 × *g* for 10 min) and resuspended in 50 mL of cold buffer A (20 mM HEPES, pH 8.0, 500 mM NaCl, 1 mM EDTA, 1 mM TCEP). This was followed by sonicated for 30 min (1 min cycles) and centrifugation (20,000 × *g* for 15 min). The expressed protein was purified by a chitin resin. Briefly, after centrifugation, the supernatant was incubated overnight with 20 mL of chitin resin in buffer A, then washed with 50 ml of buffer A and cleavage by the addition of 100 mM DTT. The resin was incubated for 48 h at room temperature on a rocking bed. The eluted solution from the chitin column was acidified to pH ~ 2.5 with 7% HCl and concentrated using Amicon Ultra-4 Centrifugal Filter Units with a membrane cut-off of 3 kDa (Merck Millipore, USA). The protein was then purified on a size exclusion chromatography SEC-70 gel filtration column (Bio-Rad) equilibrated with 10 mM HCl, then obtaining metal-free protein (apoMT2).[1] The identity of the eluted protein from SEC was confirmed by ESI-MS using a Bruker Maxis Impact (Bruker Daltonik GmbH, Bremen, Germany) calibrated with a commercial ESI-TOF Tuning mix (Sigma-Aldrich). Thiol concentration was determined spectrophotometrically using a DTNB assay[2], and the $Zn^{2+}$ binding capacity was confirmed spectrophotometrically by $Zn^{2+}$ and $Cd^{2+}$ titrations.[3] To the collected fraction of purified thionein-2, 10 molar excess of $ZnSO_4$ was added under a nitrogen blanket, and the pH was adjusted to 8.6 with a 1 M Tris base. Samples were concentrated with Amicon Ultra-4 Centrifugal Filter Units with a membrane cut-off of *3* kDa (Merck Millipore, USA) and subsequently purified on an SEC-70 gel filtration column (Bio-Rad) equilibrated with 20 mM Tris-HCl buffer at pH 8.6. Concentrations of thiols and $Zn^{2+}$ were determined spectrophotometrically using DTNB and PAR assays, respectively.[4]

**Mass spectrometry**

**Nanoelectrospray ionization.** All samples were prepared at 10-20 µM in 200 or 50 mM ammonium acetate (AmAc), pH 6.8 supplemented with 1 mM TCEP, and desalted using micro Bio-Spin 6 columns (Bio-Rad) prior to any experiment. The samples were then ionized from a borosilicate glass capillary (O.D. 1.2 mm, I.D. 0.9 mm, World Precision Instruments, Stevenage, UK) produced in-house using a Flaming/Brown P-1000 micropipette puller (Sutter Instrument Co., Novato, CA, USA). Ions were produced by applying a positive potential of 0.9-1.4 kV via a platinum wire (Goodfellow).

**Linear travelling wave ion mobility mass spectrometry.** Linear TW IM-MS experiments were carried out on a Synapt XS HDMS (Waters Corporation, Manchester, UK). Gentle source conditions were applied to prevent ion activation (source temperature 30ºC, cone voltage 20 V, source offset 1). All of the experiments were carried out in sensitivity mode to maximize ion transmission at the expense of peak resolution. Collision-induced unfolding (CIU) experiments were performed by mass-selection of ions of interest by the quadrupole mass analyzer and then by recording ion arrival time distributions under different trap collision energies (CE) in the 0-60 V range. Wave velocity and wave heights wave velocity and height were set up at 300 $ms^{-1}$ and 20 V, respectively. The helium cell and nitrogen traveling wave were operated at 200 and 75 mL·min$^{-1}$. To minimize ion activation, the trap DC bias was set up at 35 V. The mass spectra were calibrated using 2 µg·µL$^{-1}$ NaI made up of 1:1 water:isopropanol. Ion activation energies were reported as laboratory frame energy ($E_{lab}$), accounting for the charge state of the mass-selected ion. Arrival time distributions were calibrated to $^{TW}CCS_{N2}$ using a TWIMS calibration procedure. Ubiquitin (bovine), cytochrome C (equine heart), and β-lactoglobulin (bovine milk) were purchased from Sigma-Aldrich and used as calibrants. The lyophilized powders were dissolved in either 200 or 50 mM AmAc and diluted to a 10 µM protein concentration. IM-MS data was recorded on three different days, and data were averaged. The literature $CCS_{N2}$ values for the standards were obtained from A. P. France *et al.*[5]

**Optimization of salt concentration for ESI-MS studies**. Our calibration was based on protein calibrants dissolved in similar solutions conditions as for the subsequent analysis (50 mM AmAc, 1 mM TCEP). In contrast, Russell and coworkers reported a TW calibration based on denaturing calibrants.[6] Moreover, no differences in the CCSD were obtained when spraying the proteins at higher salt content, 200 mM AmAc (Figure S1). In addition, calibration of the TW cell with different wave heights and velocities yielded similar CCS values. To note, similar wave height and velocity as it was used in their studies were used here. Therefore, the CCS differences could be more likely attributed to these calibration differences.

Avoiding drastic pH drop is especially relevant for metallothioneins, where 20 thiolates coordinating $Zn^{2+}$ ions can be easily protonated, resulting in $Zn^{2+}$ dissociation. However, ammonium acetate solution does not constitute a buffer at neutral pH since the buffering properties of ammonium acetate are at pH 4.75 ± 1 and 9.25 ± 1, which corresponds to the acetic acid and ammonium p$K_a$s.[7] During the desolvation process in the ESI plume, protonation of acetate generates acetic acid (in ESI positive mode) and favors the formation of a $[M + zH]^{z+}$ ions. While this process leads to inevitable acidification, likely close to the p$K_a$ values of acetic acid, the higher the AmAc concentration, the lower the pH is shifted. For instance, a pH drop to 6.5 can be estimated when using 100 mM ammonium acetate as a solution.[7] Lowering the pH would, in turn, protonate Cys residues directly affecting the stability of the $Zn^{2+}$-binding sites.[8] Therefore, to mitigate the acidification process, the following experiments were mostly performed with 200 mM AmAc.

**Cyclic travelling wave ion mobility mass spectrometry.** Cyclic TW IMS experiments were performed on a Select Series Cyclic IMS instrument (Waters. Corporation, Manchester, UK). Samples were analyzed under similar source conditions as the Synapt XS. The TWIMS wave velocity and height were set up at 375 $ms^{-1}$ and 20 V, respectively. The helium cell and IMS nitrogen gas flow rates were 150 and 45 $mL \cdot min^{-1}$.

*Multipass cIM*. Quadrupole-selected $Zn_7MT2^{5+}$ ions (1298 *m/z*) were subjected to 1-3 passes around the cyclic device to obtain higher mobility resolving power. The trap, cyclic IM, and transfer cell were operated as gently as possible to minimize ion activation while obtaining a reasonable transmission. The ions were injected into the trap cell with an acceleration voltage of 5 V, and the post-trap bias was set up at 25 V since we could observe ion activation at standard values (45 V). The He cell bias was kept at 20 V. The ion packet from the trap cell is then injected into the cIM cell with a pre-array gradient and pre-array bias set up at 85 V, and 70 V, array offset of 25 V, and array entrance 10 V. The ions underwent 1 to 3 passes, with the wave velocity and height as abovementioned. Mobility separated ions are ejected from array to TOF analysis. Transfer collision energy was set to 10 V, to maximize the ion transmission without ion activation. Arrival time distributions (ATDs) were extracted in the [1295:1303] *m/z* range. The ATDs were deconvoluted by means CIUSuite 2 software.[9] The following parameters were used: maximum number of components = 10; peak amplitude = 0.05; peak overlap penalty mode = relaxed; expected fwhm were obtained similarly to Deslignière *et al.*[10] Briefly, a portion of ions from the convoluted ATD was selected and ejected to the pre-store while remaining ions ejected to the TOF. Then, those ions were reinjected from the pre-store to the array, separated, and the fwhm was calculated for the ATD recorded (1.50 ms). Slicing out the convoluted ATD to calculate fwhm provides accurate initial values for the peak modeling process. After one pass, a fwhm of 1.50 ms was calculated for the isolated slice. As the fwhm scales as the $\sqrt{n}$, where $n$ is the number of passes for a peak with a single conformer, we could estimate the initial fwhm for 2 (2.12 ms) and 3 passes (2.59 ms). The resolving power was calculated as CCS/ΔCCS, where ΔCCS is the extracted fwhm of the mobility peak.

*IMS-CA-IMS.* To study the presence of different ion populations in the mass-selected $Zn_7MT2^{5+}$, we used a multistage IMS[2]. As above, we utilized the same parameters in the trap (5 V), post-trap bias (25 V) and helium cell (20 V) that minimize ion activation prior to the separation in the cIM device. A mobility-selected ion population or slice was ejected and trapped in the pre-store array, while the remaining ions were ejected to the TOF. Then, the isolated ions were reinjected from the pre-store into the array. Unfolding of the mobility-selected ions was done by increasing both the pre-array gradient and the pre-array bias while keeping the voltage difference between these two parameters the same. Activated ions were subjected to one pass around the cyclic array and then ejected to the TOF. Worth comment is the need to perform two control experiments, named background ion signal and ion aging experiments.[11-12] The first one checks if the ions observed after reinjection from the pre-store are derived only from the isolated population. The second control experiment verifies the effect of time on the protein conformation. The ions are accumulated for a prolonged period of time in the array before separation.

*CA-IMS-CA-IMS.* This mode of operation is an extension to the IMS-CA-IMS with the difference that ions are also activated in the trap cell prior to the cIM.[11-12] The trap cell was used to activate all the ions before doing the IMS-CA-IMS.

**MS data analysis**. Data were analysed by means Masslynx v4.2 (Waters Corp., UK), ORIGAMI,[13] CIUSuite 2,[9] and Python 3.5 scripts.

### Computational studies

*Model building.* The X-ray structure PDB ID 4MT2, which contained four $Cd^{2+}$ and two $Zn^{2+}$ ions, was selected as the initial structure. The initial $Zn_7MT2$ was obtained by replacing the $Cd^{2+}$ with $Zn^{2+}$ ions, and point mutations were done to match with the human MT2 sequence with the VMD mutator plugin.[14] All of the simulations were performed using the GROMACS 2018.4 software.[15] The AMBER FF19SB force field was used to model the protein, and derived parameters were used to describe cysteine-$Zn^{2+}$ interactions.[16] Because of the lack of structural X-ray of NMR models for the partially $Zn^{2+}$-loaded MT2 species, $Zn_{4\text{-}6}MT2$ structures were obtained by well-tempered parallel-bias metadynamics (WT PB-MetaD).[3] Cluster analysis of the free energy minima obtained by WT PB-MetaD was used to obtain initial conformations for our studies. In particular, for $Zn_6MT2$, two configurations were included. In the first one, four $Zn^{2+}$ are bound in the α-domain and two $Zn^{2+}$ in the β-domain ($\alpha Zn_4\beta Zn_2MT2$). In the second one, three $Zn^{2+}$ are bound in each domain ($\alpha Zn_3\beta Zn_3MT2$). For $Zn_5MT2$, one representative structure was extracted by cluster analysis from the basin obtained by WT PB-MetaD

($\alpha Zn_3\beta Zn_2MT2$). For $Zn_4MT2$, two configurations were used, $\alpha Zn_2\beta Zn_2MT2$ and $\alpha Zn_3\beta Zn_1MT2$.

*Charge state distribution.* To perform an MD simulation of $[M + zH]^{z+}$ that fully represents the experimental state(s) post nESI is non-trivial. The inclusion of water-water and water-protein proton transfer events cannot be captured by standard force fields[17] and would require the use of quantum mechanics (QM) treatments. Including ~ 2500 molecules in a QM part of a QM/MM scheme is computationally prohibitive nowadays. Here, we incorporated three alternative solutions to tackle this issue, namely: (i) a mobile $Na^+$ charge scheme in which all titratable residues were set up to their default pH 7.0 protonation state, except for the Cys residues, and $Na^+$ ions carried out the positive charges. Here, the Cys residues that had bound $Zn^{2+}$ were modeled in the deprotonated form as this is how they are found experimentally in the X-ray structure and determined in this study by MS. In the partially $Zn^{2+}$-depleted and metal-free MT2 species, the free Cys residues were modeled as protonated in agreement with our MS data; (ii) a mobile $Na^+$/static $H^+$ charge scheme in which the negative charges (Asp and Glu residues) were neutralized while protonating the positive residues (Lys residues).[18] We should consider that $Zn^{2+}$ carries a 2+ charge, each Cys-binding residue has a -1 charge, and a free Cys residue is neutral. In our case, metallothionein does not contain enough residues to distribute the *z* excess protons on the protein. In this approach, it is not fully correct to consider that the acidic sites are generally neutral since carboxylates R-$COO^-$ sites involved in salt bridges have been found.[18] Then, in all of the systems, several $Na^+$ were added to obtain a total system charge of 5+ and; (iii) a mobile $H^+$ approach that considers that the protons are highly mobile and the preferred proton-binding residues can change as the simulation, and therefore the protein structure evolves.[19-20] We used a recently released charge placement algorithm (ChargePlacer),[21] albeit modified to incorporate $Zn^{2+}$-binding residues in determining of the protonation pattern. The MD simulation was divided into multiple 50 ps NVT runs, and at the beginning of each simulation, the charge placement algorithm redistributed the protons along all of the titratable residues and all of the 20 Cys residues, independently if they had bound a $Zn^{2+}$ ion. Briefly, the ChargePlacer algorithm finds the protonation pattern that minimizes the total energy of the system, which accounts for both Coulomb repulsion and proton affinity, resembling the approach described by Konermann.[19-20]

*Gas-phase desolvation MD.* Each protein system was solvated in a rhombic dodecahedron box with ~ 2500 TIP4P/2005 water molecules, since they provide more realistic results of the ESI droplet evaporation.[18] The three-site TIP3P water molecule is treated as a nonpolarizable molecule and exhibits a lower ~30% surface tension than the real water molecules. Here we employed the $Na^+$ mobile approach by which the aqueous droplet was charged by randomly replacing water molecules with excess $Na^+$ to obtain a system total charge of 16+. According to the charge residue mechanism (CRM) the maximum charge of positive charges that an ion can obtain is the Rayleigh limit ($z_R$).[22] For globular spherical protein, $z_R = 0.0778\ m^{1/2}$, where $m$ is the molecular weight of the protein in Da units.[23] The value of $z_R$ can deviate for proteins that are not structurally globular.[24] To account for any deviation, we employed a $2.5z_R$ excess of $Na^+$ ions in the initial droplets. MD runs with $z \sim z_R$ yielded similar final $[M+ zNa]^{z+}$ ions and $z/z_R$.

We carried out a pseudo-PBC approach in which the Coulomb and Lennard-Jones cutoffs were set to 300 nm, and the PBC box dimensions were set up to 900 $nm^3$. Although this approach uses periodic boundary conditions, the box size and cutoffs exclude interactions between PBC images. Each system was subjected to energy minimization by 10 000 steps of steepest descent minimization, followed by 10 ps NVT equilibration to 350 K by using the Nosé-Hoover thermostat with a coupling constant of 0.1 ps. The LINCS algorithm was used to constraint bonds involving hydrogen atoms to be able to use a 2 fs time step, and neighbor list was updated every 100 steps using the Verlet method. To simulate droplet desolvation, the MD simulations were split into multiple consecutive 250 ps length NVT runs. At the end of each window, water molecules and $Na^+$ and/or $Zn^{2+}$ ions further than 30 Å from the center of mass of the protein were removed. The system was then recentered in the box, and the velocities reassigned from a Maxwell-Boltzmann distribution. This approach avoids evaporative cooling problems and speeds up the MD simulations as the number of particles is reduced. After 250 runs or 62.5 ns of NVT production at 350 K, the system was equilibrated to 500 K, and run for 5 ns to remove the last water molecules, also called "sticky" waters. Two independent runs were performed for each system, amounting to 0.9 µs of dynamics.

Collision cross section values were calculated every 250 ps using the trajectory method implemented in IMPACT software.[25] The simulations were analyzed using MDAnalysis 2.0[26-27], MDTraj 1.98[28], pytraj 2.0.5,[29] and in-house Python 3.5 scripts.

*Gas-phase MD simulation.* In contrast to the desolvation protocol, here, each protein system was first equilibrated in the presence of solvent molecules and then directly placed in pseudo-vacuum conditions. To compare it with our experimental data, the (i) mobile $Na^+$ charge scheme, (ii) mobile $Na^+$/static $H^+$ charge scheme, and a (iii) mobile $H^+$ approach were considered to capture the charge state distribution for the $[M + 5H]^{5+}$ ions. As above, we used a pseudo-PBC approach and ran each system for 100 ns at 298 K, and 798 K on three replicate runs for the (i) and (ii) approaches. While approaches (i) and (ii) were run for the seven protein systems, approach (iii) was run twice for 1500 ns at 298 K exclusively for the $Zn_7$MT2 structure. The cumulative simulation time approached 11.4 μs.

*Simulated Annealing.* To simulate protein unfolding as obtained by CIU experiments, structures obtained after gas-phase desolvation underwent a simulated annealing (SA) protocol with the pseudo-PBC approach. In the SA protocol the temperature was rise linearly from 298 to 798 K during 10 ns, we also compared results when applying a 100 ns run. Each system was run on three replicate runs. A total of 210 ns of SA were run.

*Steered molecular dynamics (SMD) simulations*. SMD was used to study the nonequilibrium unfolding dynamics of the protein systems obtained after gas-phase desolvation. Identical parameters as above were used for pseudo-PBC, temperature coupling, and bond parameters. The SMD computations were performed with GROMACS 2018.4 in combination with the PLUMED plugin.[30] Two different collective variables (CV) were considered: the end-to-end Met1(CA)-Ala61(CA) distance and the radius of gyration ($R_g$). In the first CV, constant force of 95 kcal·mol$^{-1}$·nm$^{-2}$ with a pulling speed of 10 Å·ns$^{-1}$ was used to produce extensions up to 50 Å so that $Zn^{2+}$ dissociation does not occur, as observed during collision-induced unfolding experiments. The pulling force was optimized to be high enough to observe a linear dependence between the distance and the CV position while not producing high fluctuations in the force profile. Three force constants of 48, 95, and 190 kcal·mol$^{-1}$·nm$^{-2}$ were tested. Using a constant force of 95 kcal·mol$^{-1}$·nm$^{-2}$ was considered optimum as it accomplished the abovementioned requirements but also maintained the native number of Zn-S bonds in the structure and thus kept the $Zn^{2+}$ binding sites in their most native state as observed experimentally. The relationship between the pulling speed and rupture or unfolding force was also considered by using several pulling rates: 1, 10, and 100 Å·ns$^{-1}$. The simulation time needed for each pulling speed was calculated as $(r_F\text{-}r_0)/k$, where $r_F$ is the final distance, $r_0$ is the initial distance, and k is the pulling speed. Thus, 50, 5, or 0.5 ns were run for each pulling rate assayed, respectively. A pulling speed of 10 Å·ns$^{-1}$ was chosen as it gave comparable unfolding forces as the lowest speed considered (1 Å·ns$^{-1}$) but saved computational time. In another set of SMD simulations, *R*g was employed as a CV. Initials trials established the maximum *R*g value needed to promote a conformational transition from a compact to an unfolded structure with intact $Zn^{2+}$ binding sites. Thus, *R*g moved from an initial value of 11 Å to 18 Å. As above, three different force constants of 10, 25, and 50 kcal·mol$^{-1}$·nm$^{-2}$ were tested. The most suitable force constant was 25 kcal·mol$^{-1}$·nm$^{-2}$ with a pulling speed of 10 Å·ns$^{-1}$ as it provided a linear relationship between the distance and position of the CV and did not induce $Zn^{2+}$ dissociation events. In total, 225 ns of SMD simulation time was considered for analysis. The RMSD, number of Zn–S bonds, number of hydrogen bonds (h-bonds), and number of salt bridges were computed by MDAnalysis 2.0[26-27] and Python 3.5 scripts. Statistical analysis of the forces and work were performed by using a two-tailed *t*-test with equal variances.

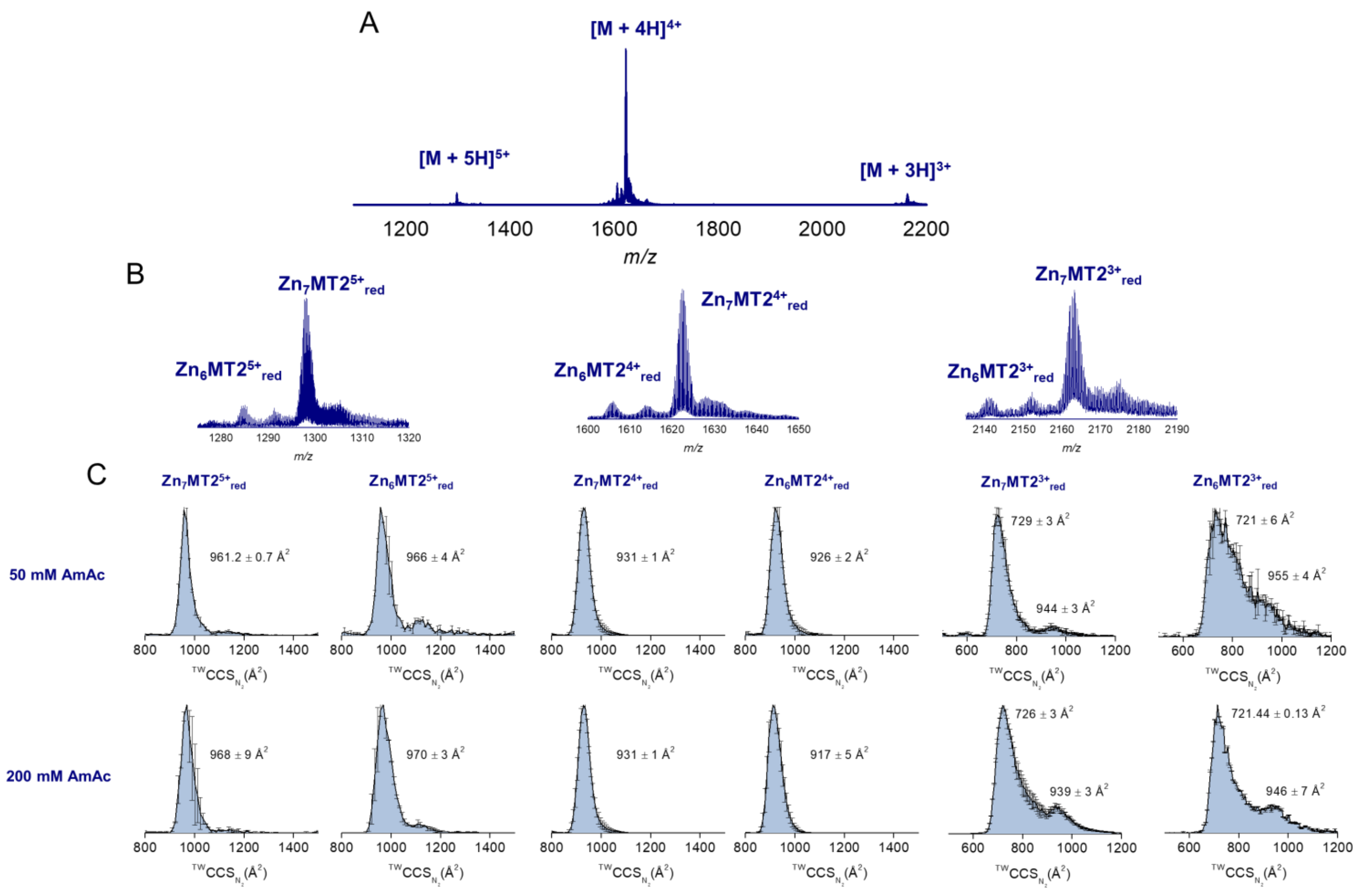


**Figure S1.** Native MS and CCS distributions for $Zn_7MT2$ (10 µM) sprayed under 50 and 200 mM AmAc supplemented with 1 mM TCEP. The CCS values were calculated from three replicates, and the error bars plot along the CCS axis.

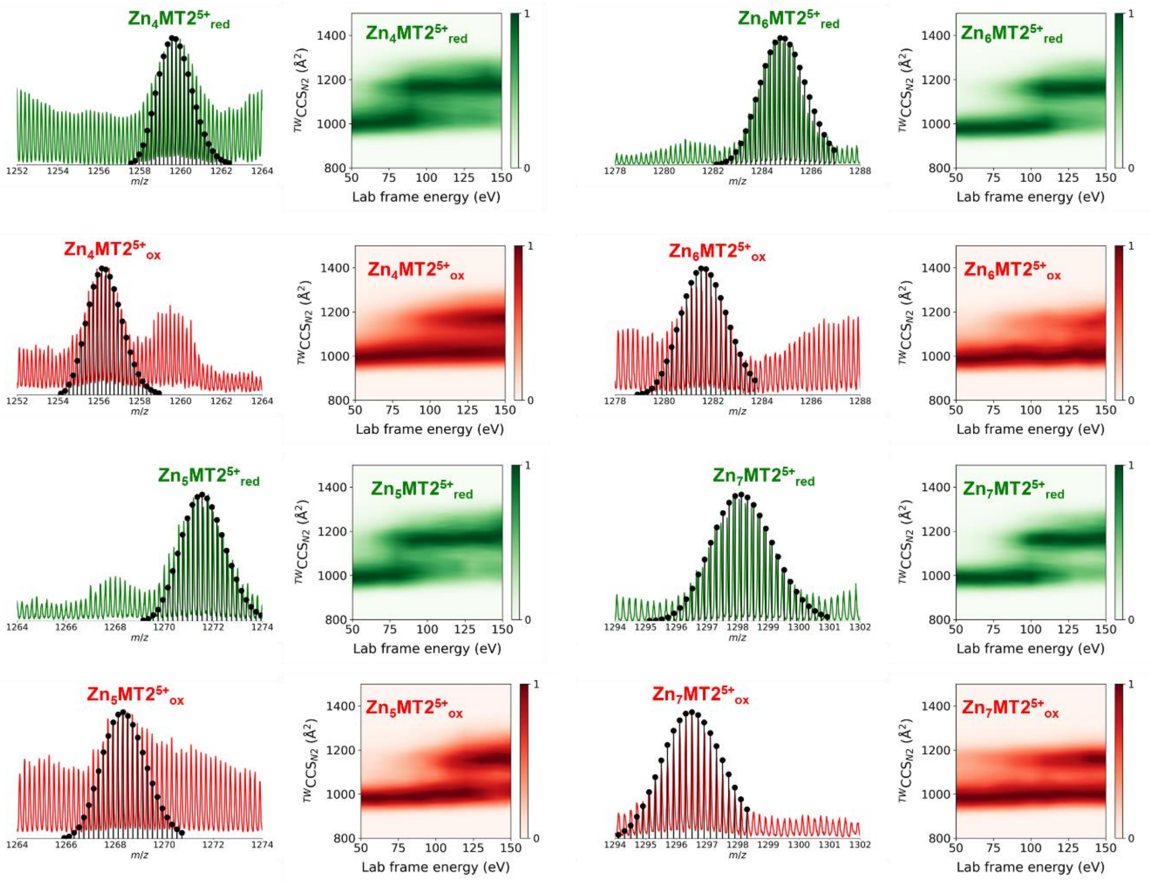


**Figure S2.** Native mass spectra and collision-induced unfolding (CIU) heat maps for the mass-selected 5+ ions of reduced and oxidized $Zn_xMT2$ (x = 4-7) sprayed from 50 mM ammonium acetate (pH 6.8) in the presence (reduced forms, “red”) and absence (oxidized, “ox”) of 1 mM pH neutralized TCEP (pH 7.4). Theoretical isotopic patterns are shown in black as stem plots, and the molecular formulas are shown in **Table S1**. Activation of the ions was performed in the trap cell prior to the IM cell by applying a linear collision energy ramp between 5-50 V with increments of 5 V.

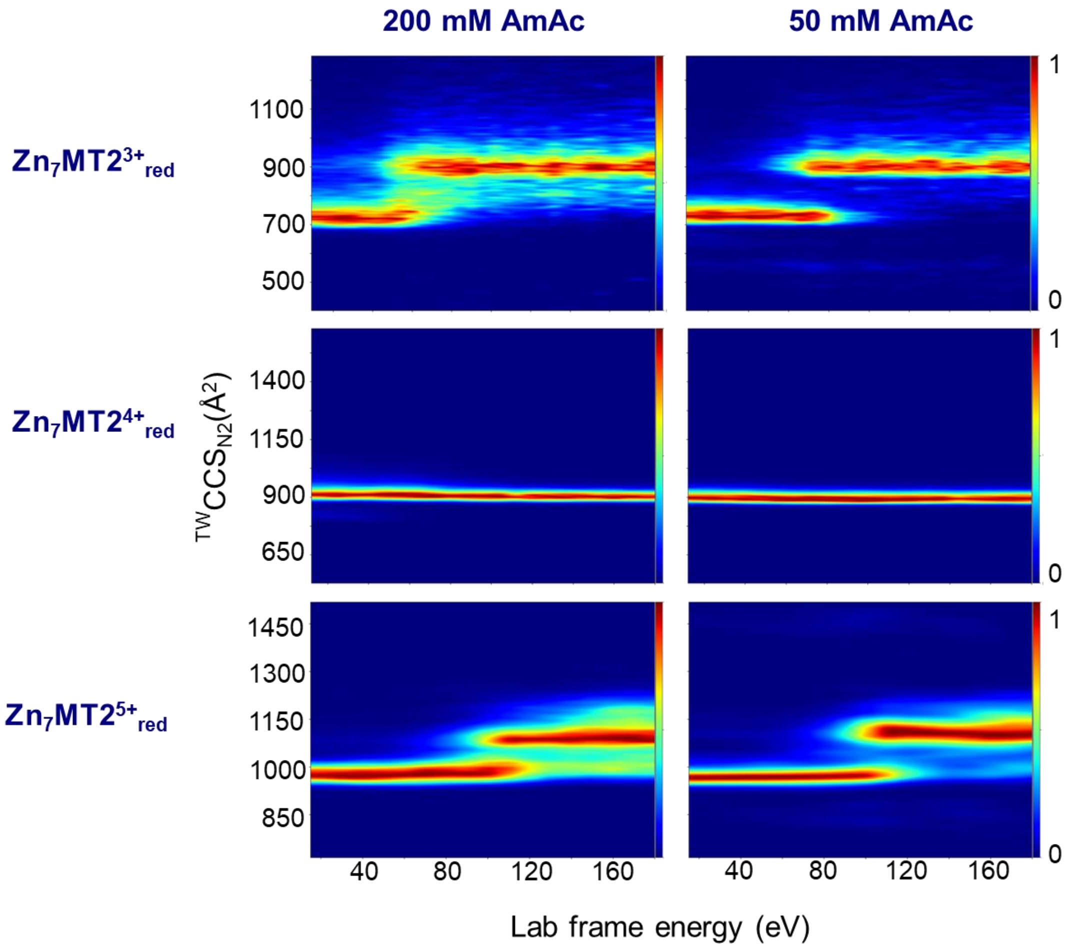


**Figure S3.** Collision induced unfolding (CIU) heat maps for $Zn_7MT2^{Z+}{}_{red}$ ($z$ = 3-5) sprayed from 50 or 200 mM ammonium acetate (AmAc) supplemented with 1 mM TCEP.

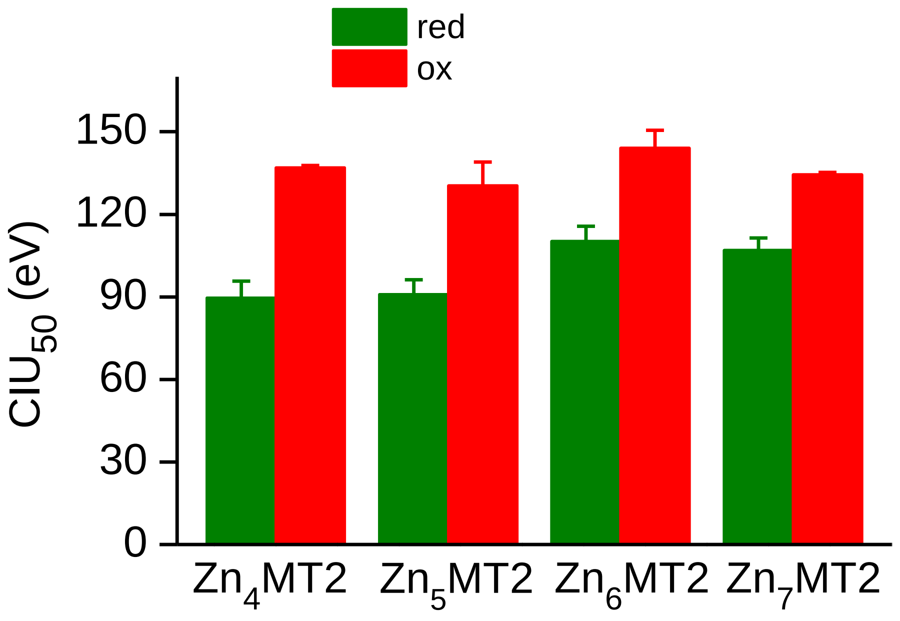


**Figure S4.** Gas-phase stabilities of $Zn_{4\text{-}7}MT2^{5+}$ ions measured as $CIU_{50}$, which in our case refers to the energy required to promote a conformational transition of 50 % of the ions from a compact to an extended conformation. $CIU_{50}$ was calculated by fitting the collision-cross section distributions along the collision energies applied. Red and ox refer to reduced and oxidized ions, respectively.

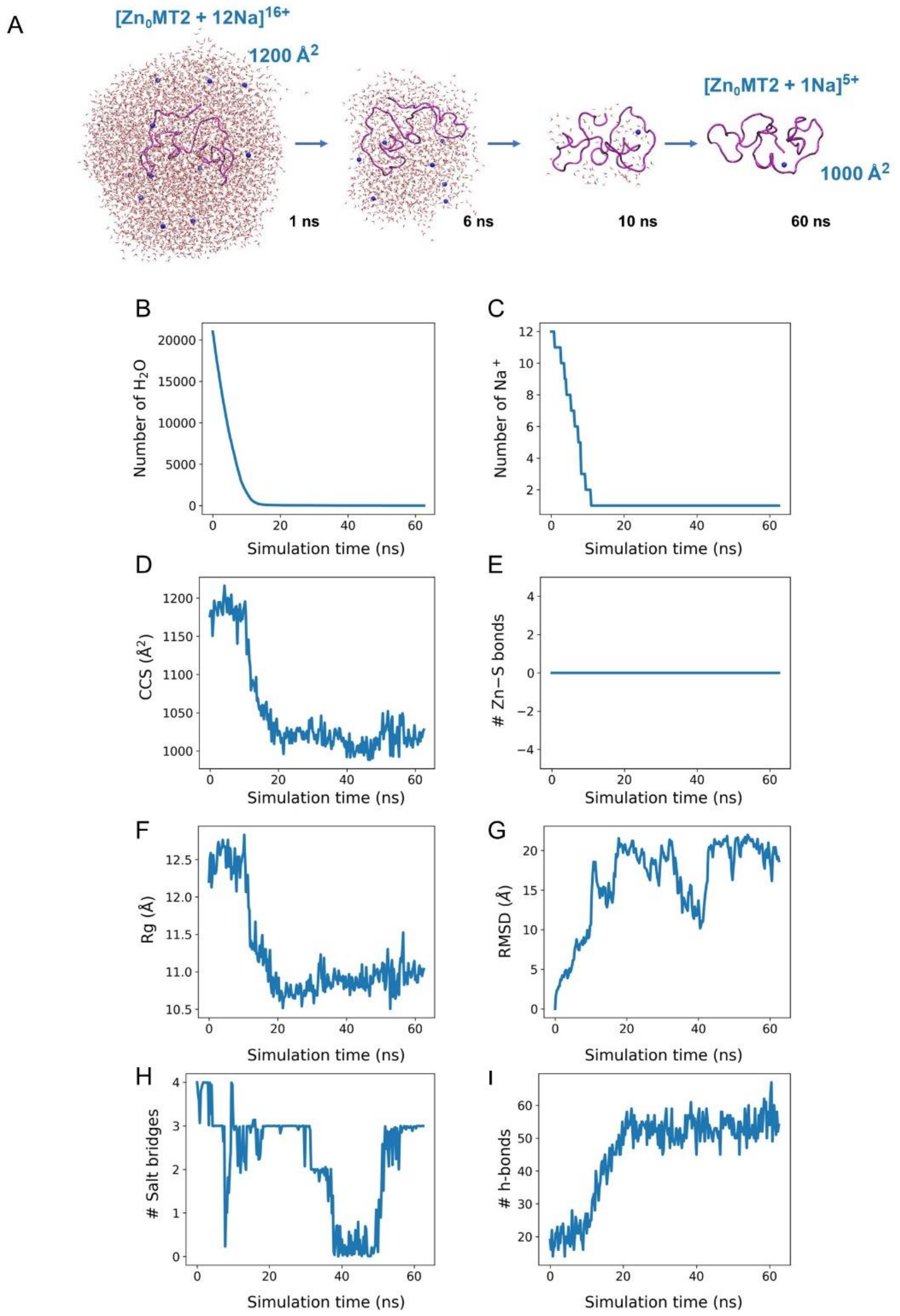


**Figure S5.** Gas-phase MD simulations of the electrospray ionization process of aqueous nanodroplet containing $Zn_0$MT2 and $Na^+$ to achieve a 16+ overall charge. (A) Snapshots of the desolvation process at different frames. $Na^+$ is represented by a blue sphere, the protein backbone is shown in magenta, and the oxygen atoms from solvent molecules are shown in red. The number of water molecules (B), Na+ ions (C), CCS values (D), number of Zn–S bonds (E), the radius of gyration (F), root-mean-square deviation (G), number of salt bridges (H) and hydrogen bonds (I) were monitored throughout the desolvation.

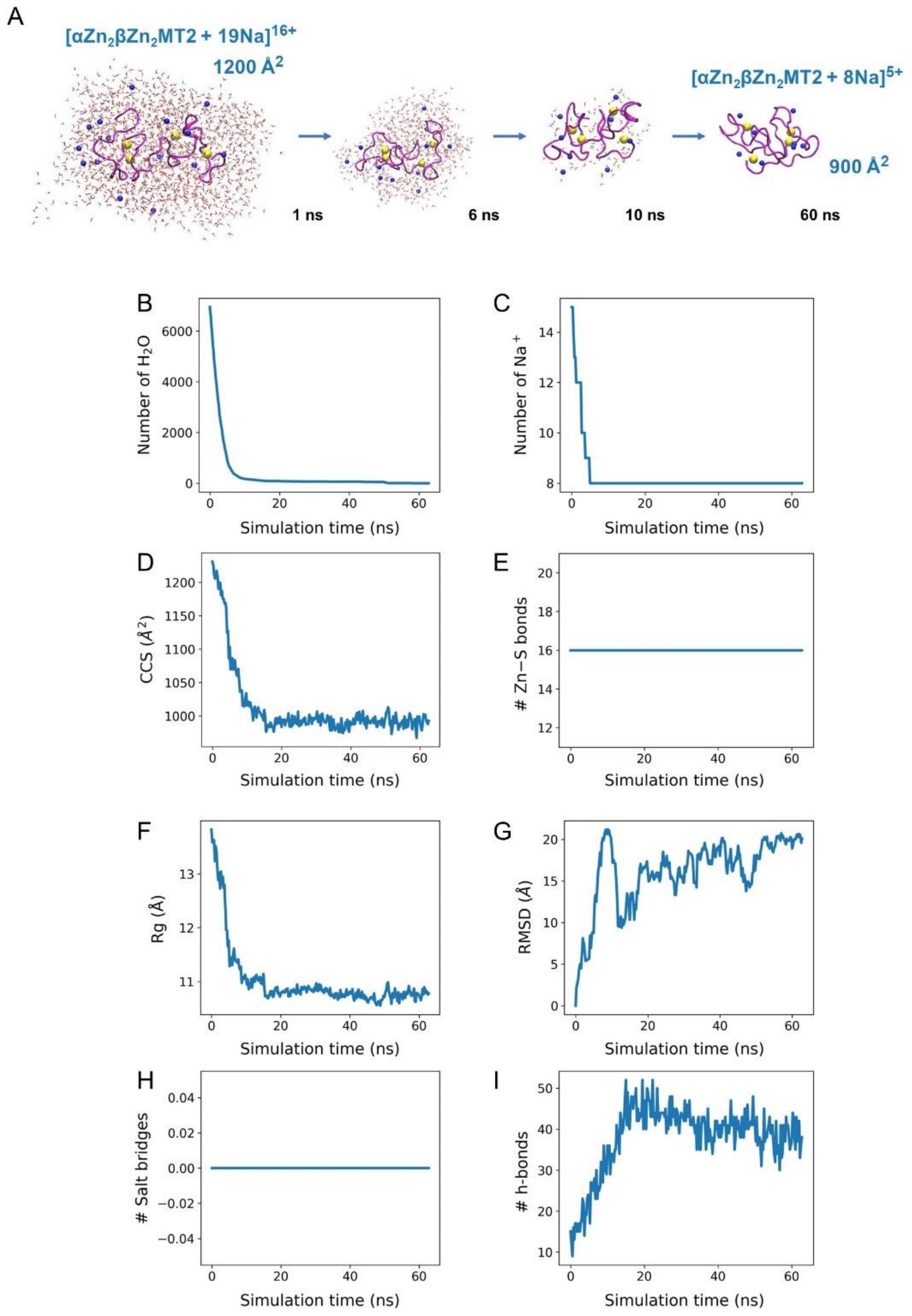


**Figure S6.** Gas-phase MD simulations of the electrospray ionization process of aqueous nanodroplet containing $\alpha Zn_2\beta Zn_2MT2$ and $Na^+$ to achieve a 16+ overall charge. (A) Snapshots of the desolvation process at different frames. $Na^+$ is represented by a blue sphere, the protein backbone is shown in magenta, and the oxygen atoms from solvent molecules are shown in red. The number of water molecules (B), $Na^+$ ions (C), CCS values (D), number of Zn−S bonds (E), the radius of gyration (F), root-mean-square deviation (G), number of salt bridges (H) and hydrogen bonds (I) were monitored throughout the desolvation.

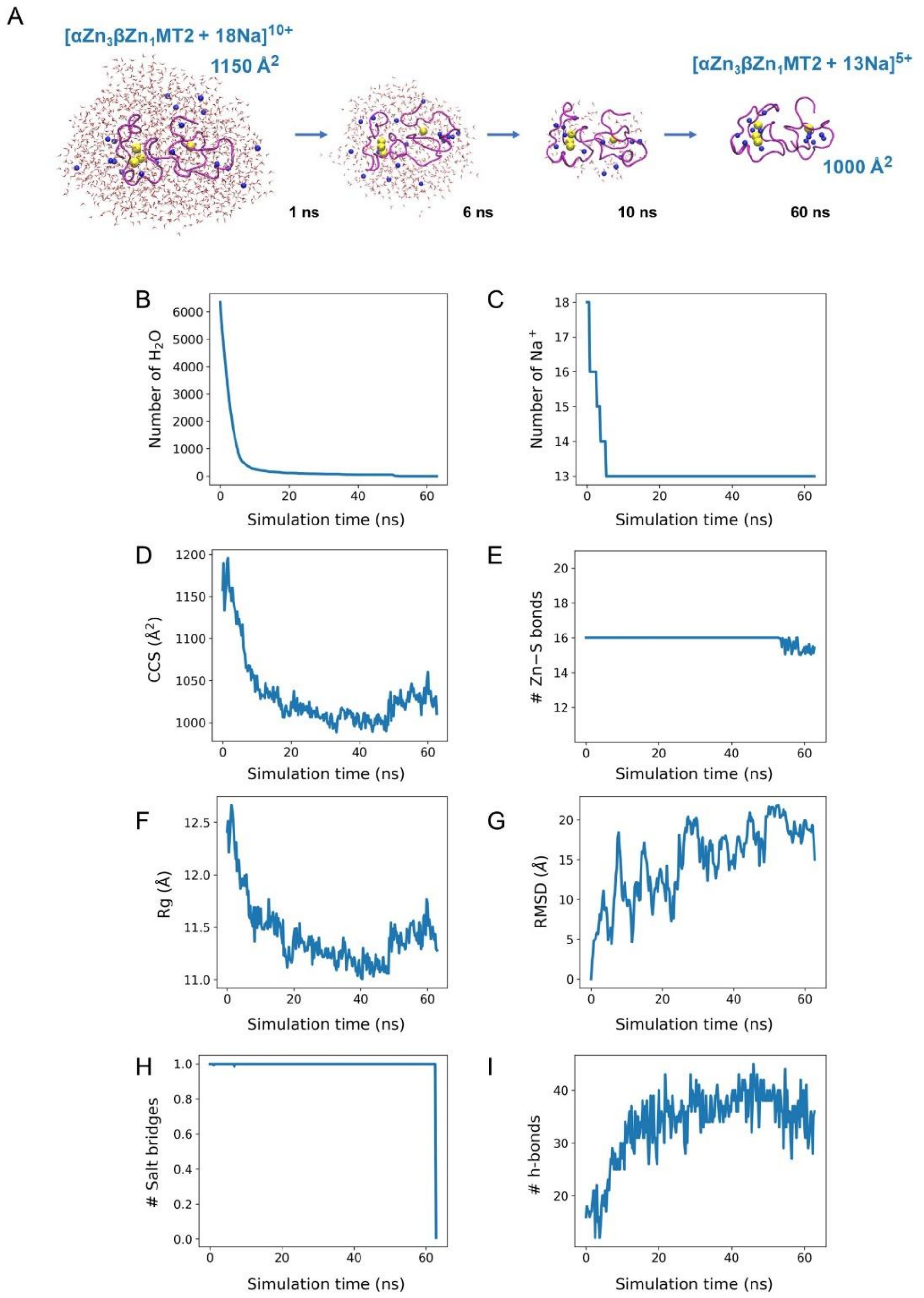


**Figure S7.** Gas-phase MD simulations of the electrospray ionization process of aqueous nanodroplet containing $\alpha Zn_3\beta Zn_1$MT2 and $Na^+$ to achieve a 16+ overall charge. (A) Snapshots of the desolvation process at different frames. $Na^+$ is represented by a blue sphere, the protein backbone is shown in magenta, and the oxygen atoms from solvent molecules are shown in red. The number of water molecules (B), $Na^+$ ions (C), CCS values (D), number of Zn−S bonds (E), the radius of gyration (F), root-mean-square deviation (G), number of salt bridges (H) and hydrogen bonds (I) were monitored throughout the desolvation.

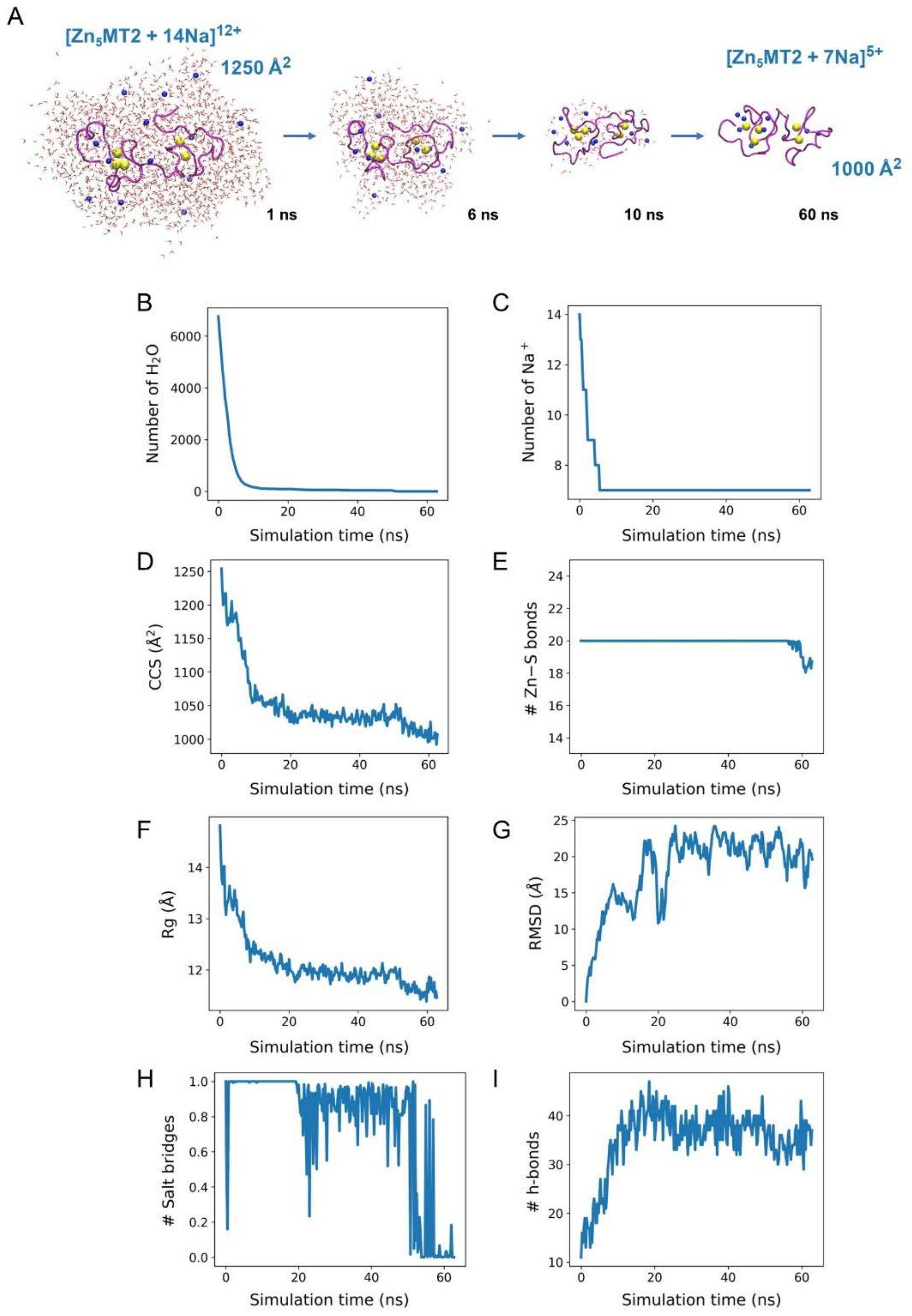


**Figure S8.** Gas-phase MD simulations of the electrospray ionization process of aqueous nanodroplet containing $Zn_5$MT2 and $Na^+$ to achieve a 16+ overall charge. (A) Snapshots of the desolvation process at different frames. $Na^+$ is represented by a blue sphere, the protein backbone is shown in magenta, and the oxygen atoms from solvent molecules are shown in red. The number of water molecules (B), $Na^+$ ions (C), CCS values (D), number of Zn─S bonds (E), the radius of gyration (F), root-mean-square deviation (G), number of salt bridges (H) and hydrogen bonds (I) were monitored throughout the desolvation.

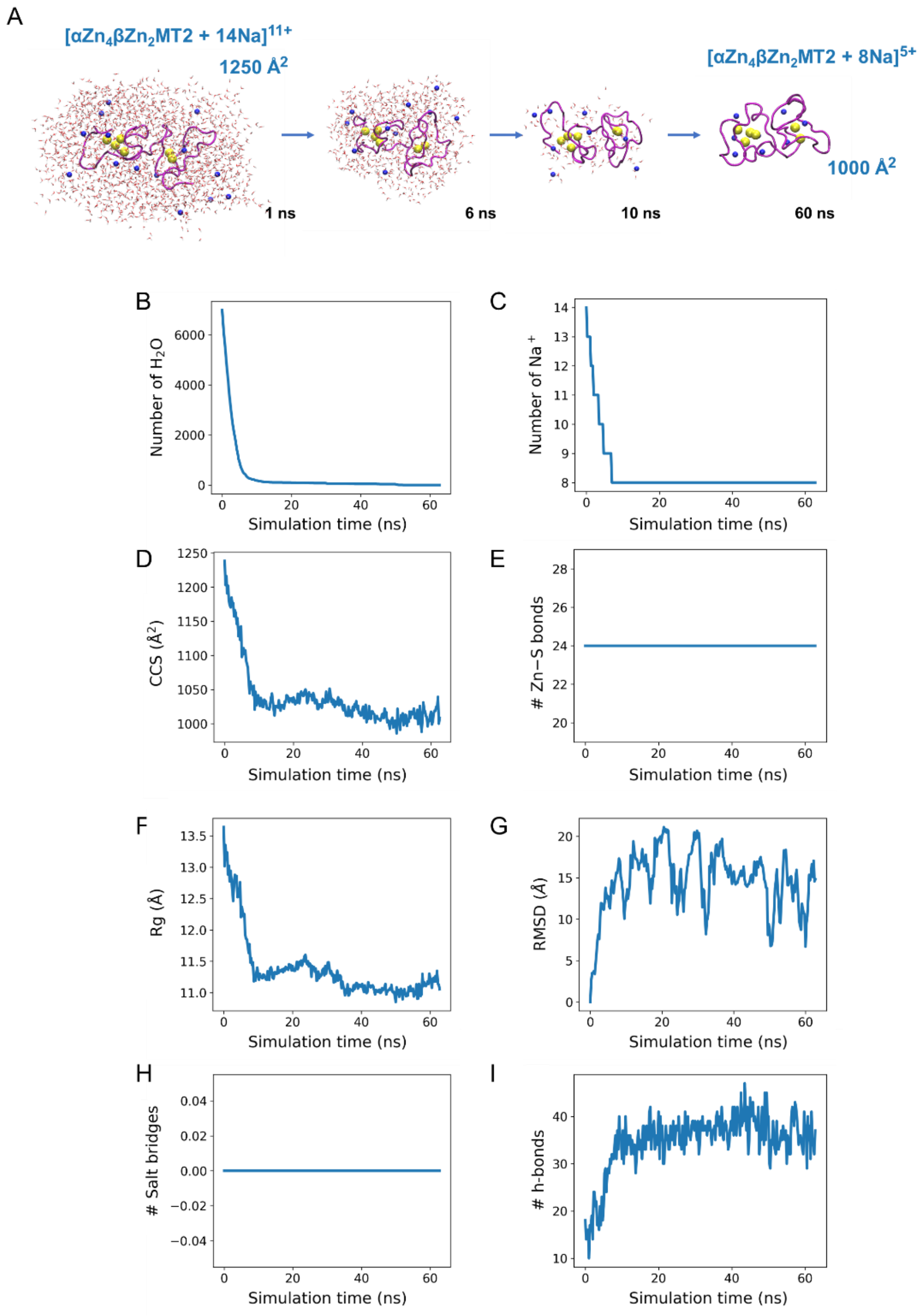


**Figure S9.** Gas-phase MD simulations of the electrospray ionization process of aqueous nanodroplet containing $\alpha Zn_4\beta Zn_2MT2$ and $Na^+$ to achieve a 16+ overall charge. (A) Snapshots of the desolvation process at different frames. $Na^+$ is represented by a blue sphere, the protein backbone is shown in magenta, and the oxygen atoms from solvent molecules are shown in red. The number of water molecules (B), $Na^+$ ions (C), CCS values (D), number of Zn−S bonds (E), the radius of gyration (F), root-mean-square deviation (G), number of salt bridges (H) and hydrogen bonds (I) were monitored throughout the desolvation.

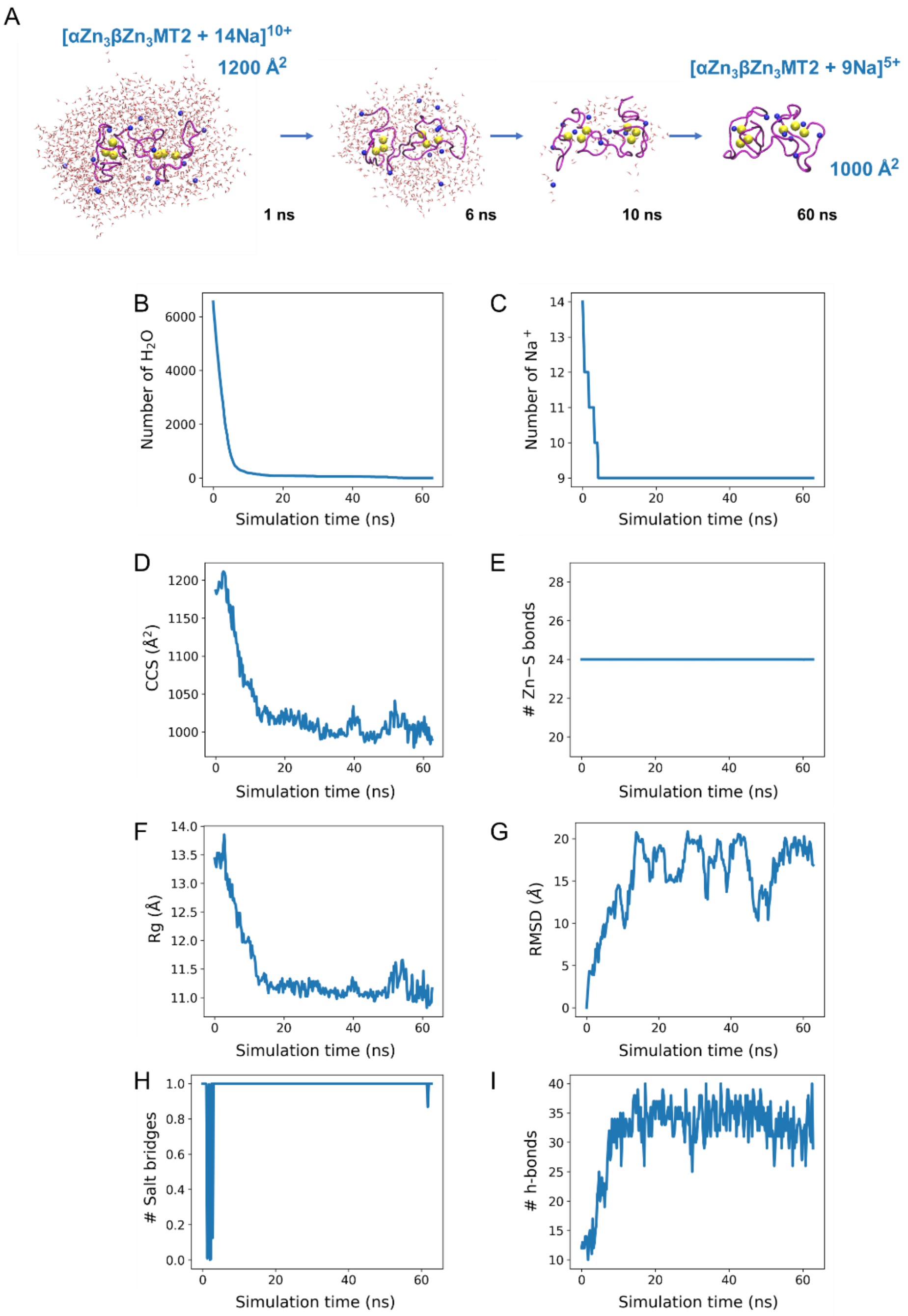


**Figure S10.** Gas-phase MD simulations of the electrospray ionization process of aqueous nanodroplet containing αZn3βZn3MT2 and $Na^+$ to achieve a 16+ overall charge. (A) Snapshots of the desolvation process at different frames. $Na^+$ is represented by a blue sphere, the protein backbone is shown in magenta, and the oxygen atoms from solvent molecules are shown in red. The number of water molecules (B), $Na^+$ ions (C), CCS values (D), number of Zn−S bonds (E), the radius of gyration (F), root-mean-square deviation (G), number of salt bridges (H) and hydrogen bonds (I) were monitored throughout the desolvation.

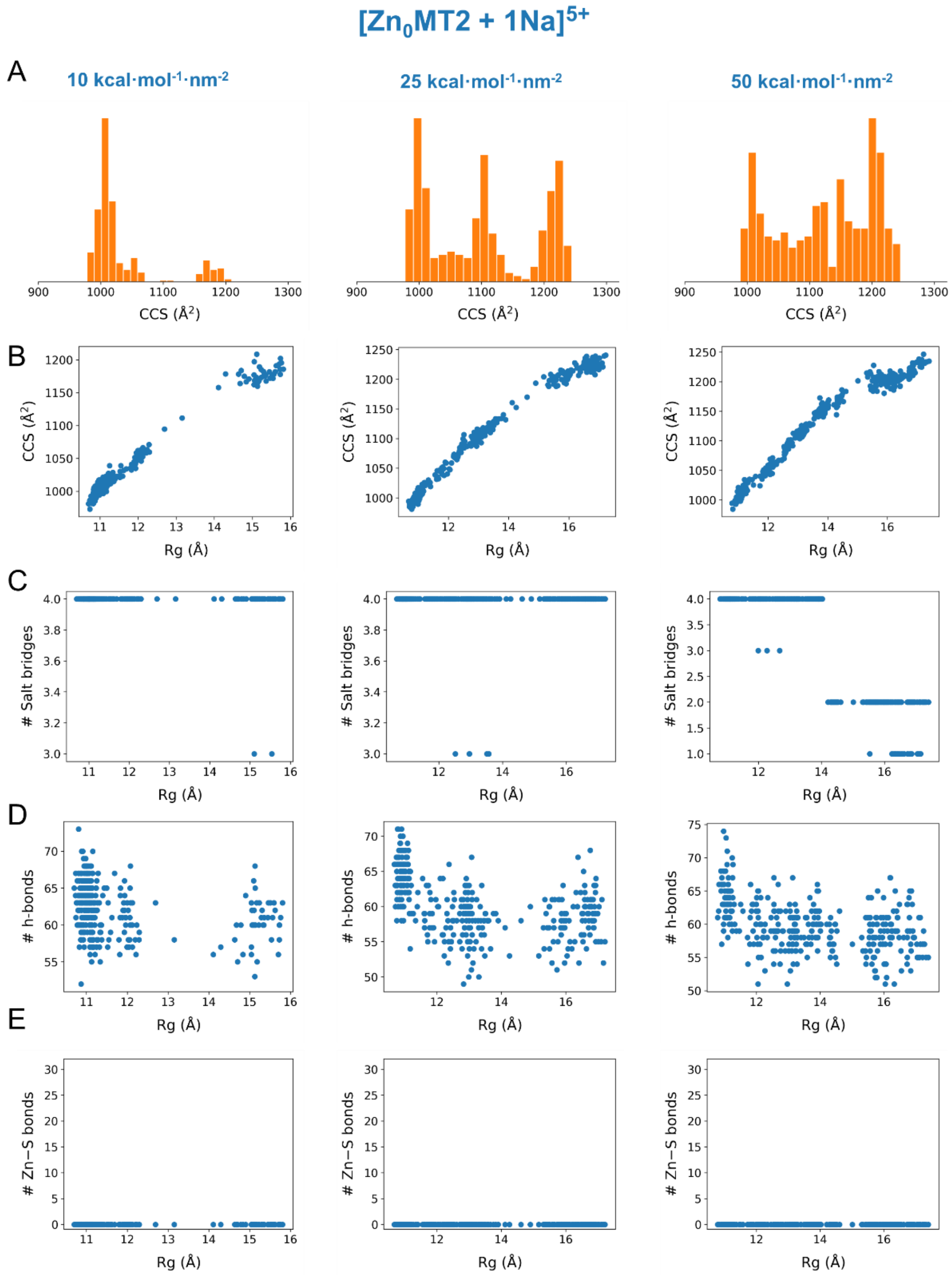


**Figure S11.** Optimization of the SMD simulations involving gas-phase $[Zn_0MT2 + 1Na]^{5+}$ ions using the radius of gyration ($R_g$) as a collective variable. CCS histograms for three different force constants (10, 25, and 50 kcal·mol$^{-1}$·nm$^{-2}$) (A), CCS (B), number of salt bridges (C), number of hydrogen bonds (D) and number of Zn−S bonds (E) as a function of the $R_g$.

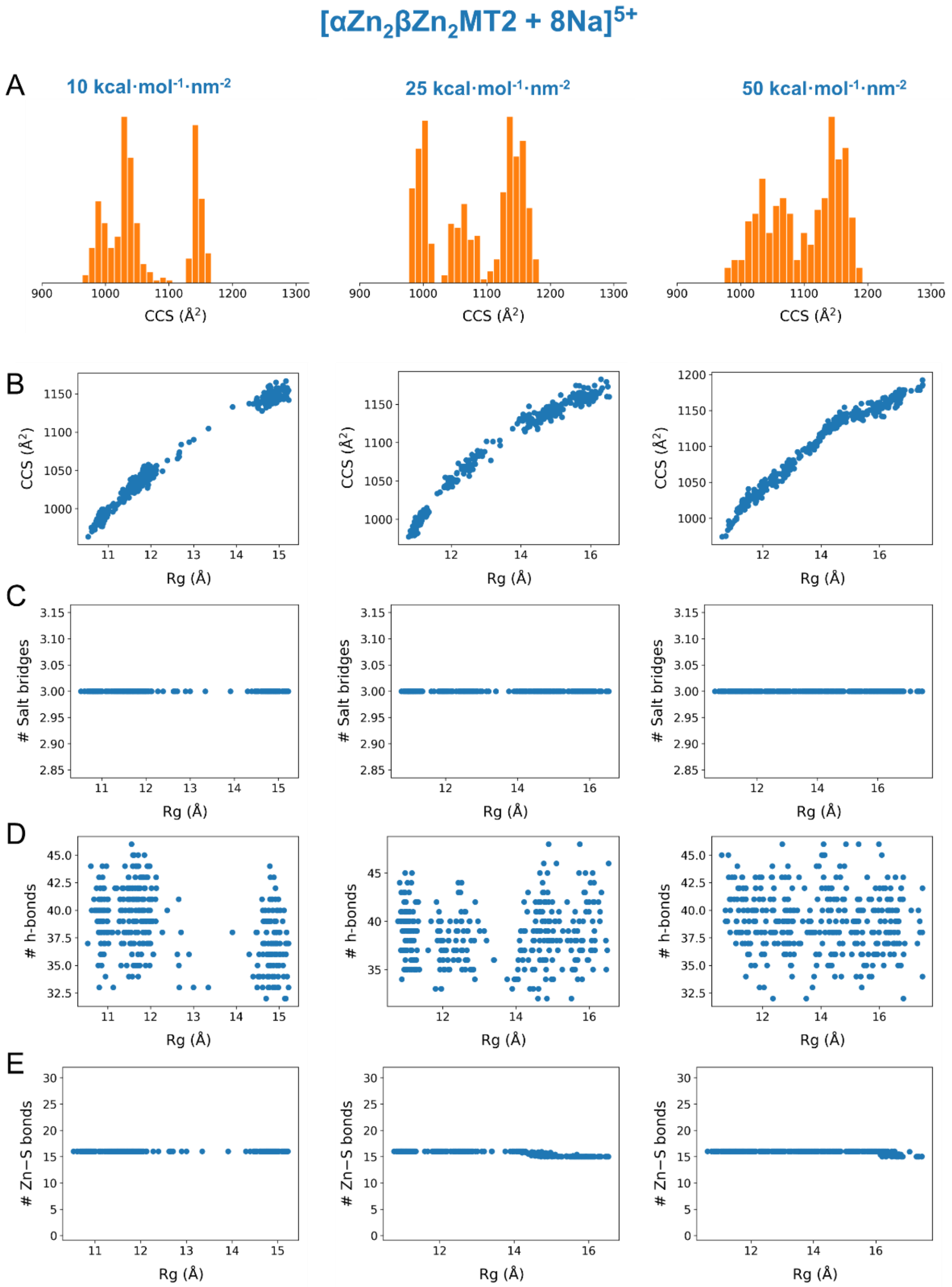


**Figure S12.** Optimization of the SMD simulations involving gas-phase [$\alpha Zn_2\beta Zn_2$MT2 +8Na]$^{5+}$ ions using radius of gyration ($R_g$) as a collective variable. CCS histograms for three different force constants (10, 25, and 50 kcal·mol$^{-1}$·nm$^{-2}$) (A), CCS (B), number of salt bridges (C), number of hydrogen bonds (D) and number of Zn−S bonds (E) as a function of the $R_g$.

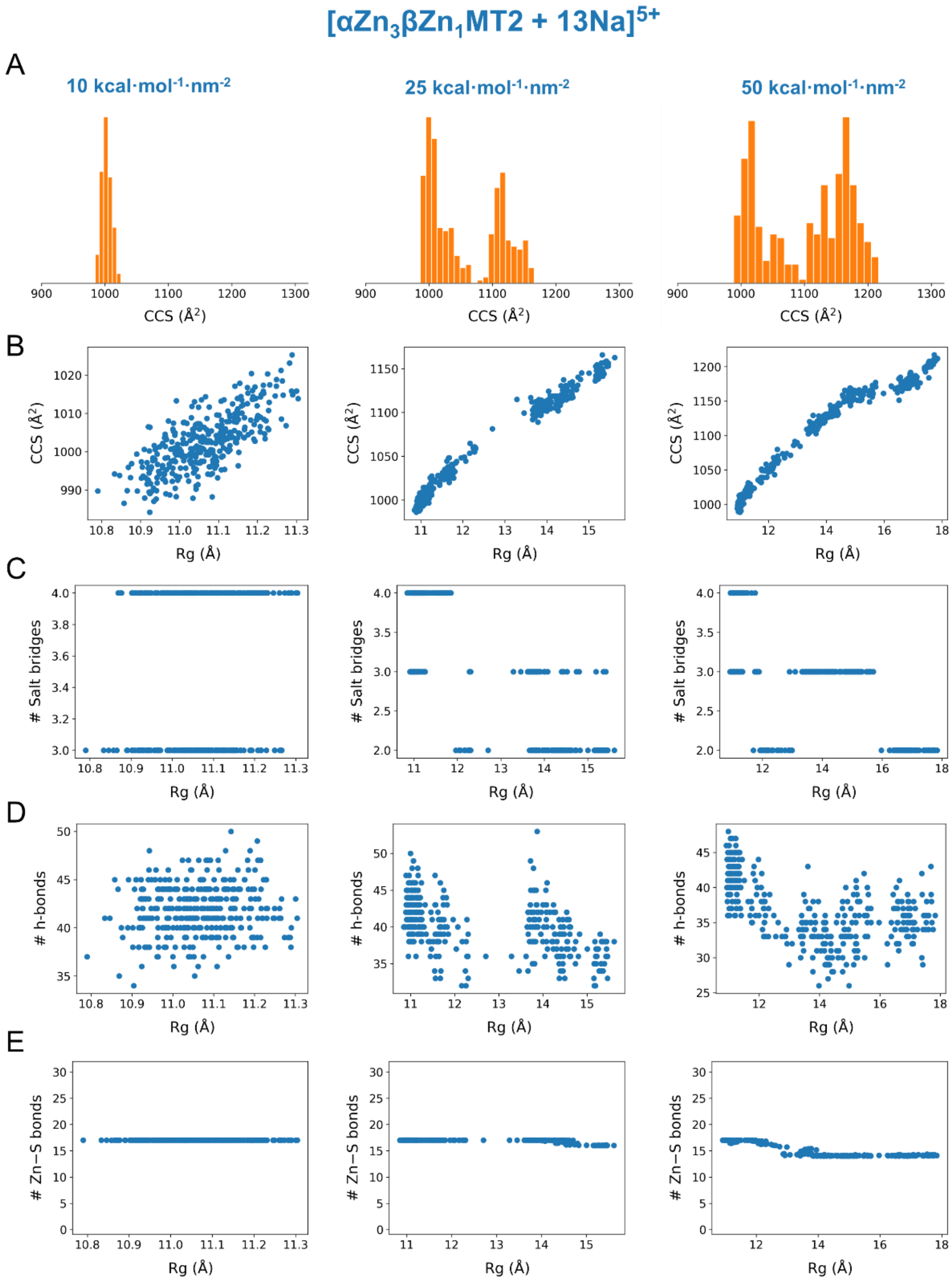


**Figure S13.** Optimization of the SMD simulations involving gas-phase [$\alpha Zn_3\beta Zn_1$MT2 +13Na]$^{5+}$ ions using radius of gyration ($R_g$) as a collective variable. CCS histograms for three different force constants (10, 25, and 50 kcal·mol$^{-1}$·nm$^{-2}$) (A), CCS (B), number of salt bridges (C), number of hydrogen bonds (D) and number of Zn−S bonds (E) as a function of the $R_g$.

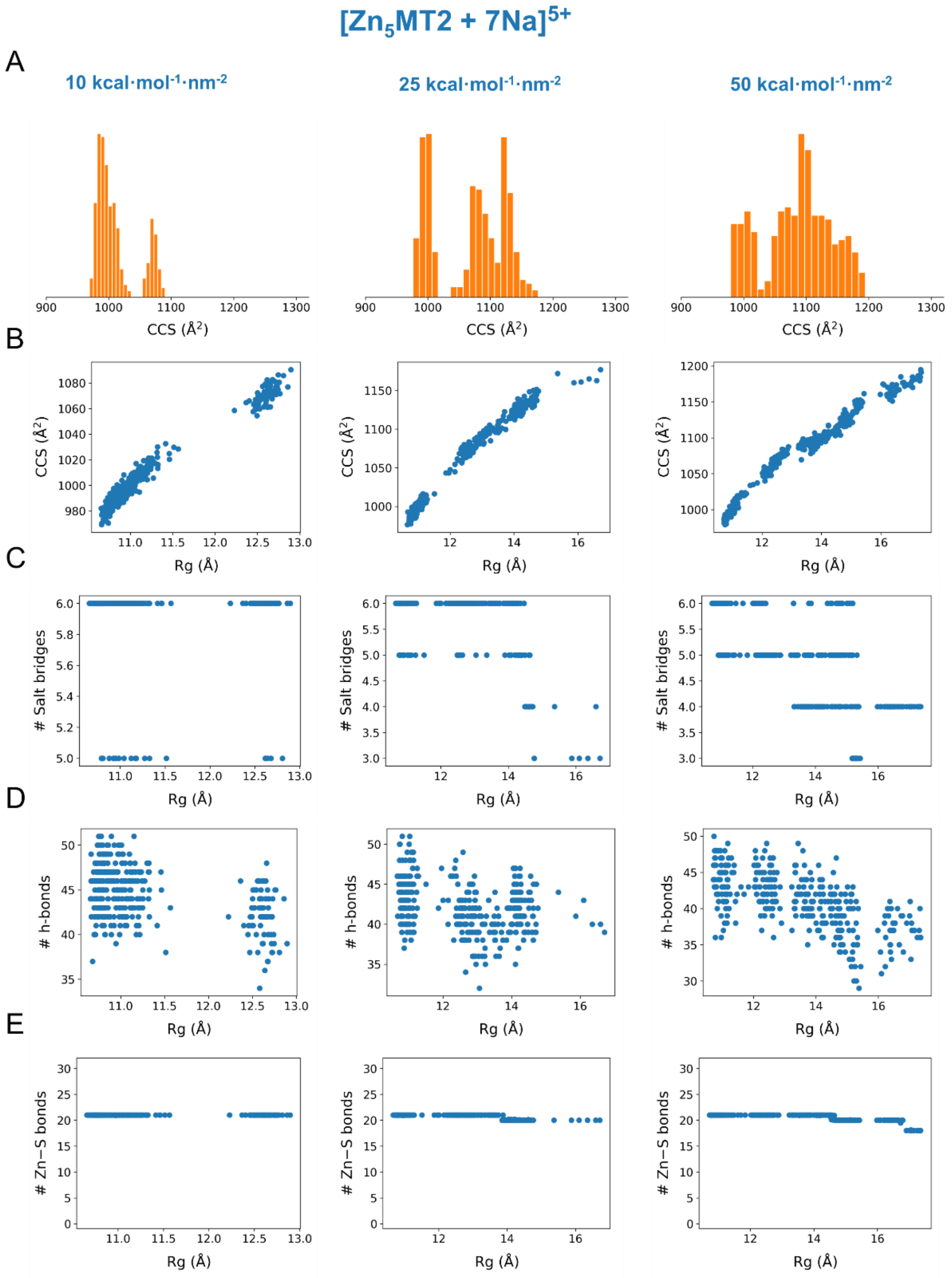


**Figure S14.** Optimization of the SMD simulations involving gas-phase [$Zn_5$MT2 +7Na]$^{5+}$ ions using radius of gyration ($R_g$) as a collective variable. CCS histograms for three different force constants (10, 25, and 50 kcal·mol$^{-1}$·nm$^{-2}$) (A), CCS (B), number of salt bridges (C), number of hydrogen bonds (D) and number of Zn−S bonds (E) as a function of the $R_g$.

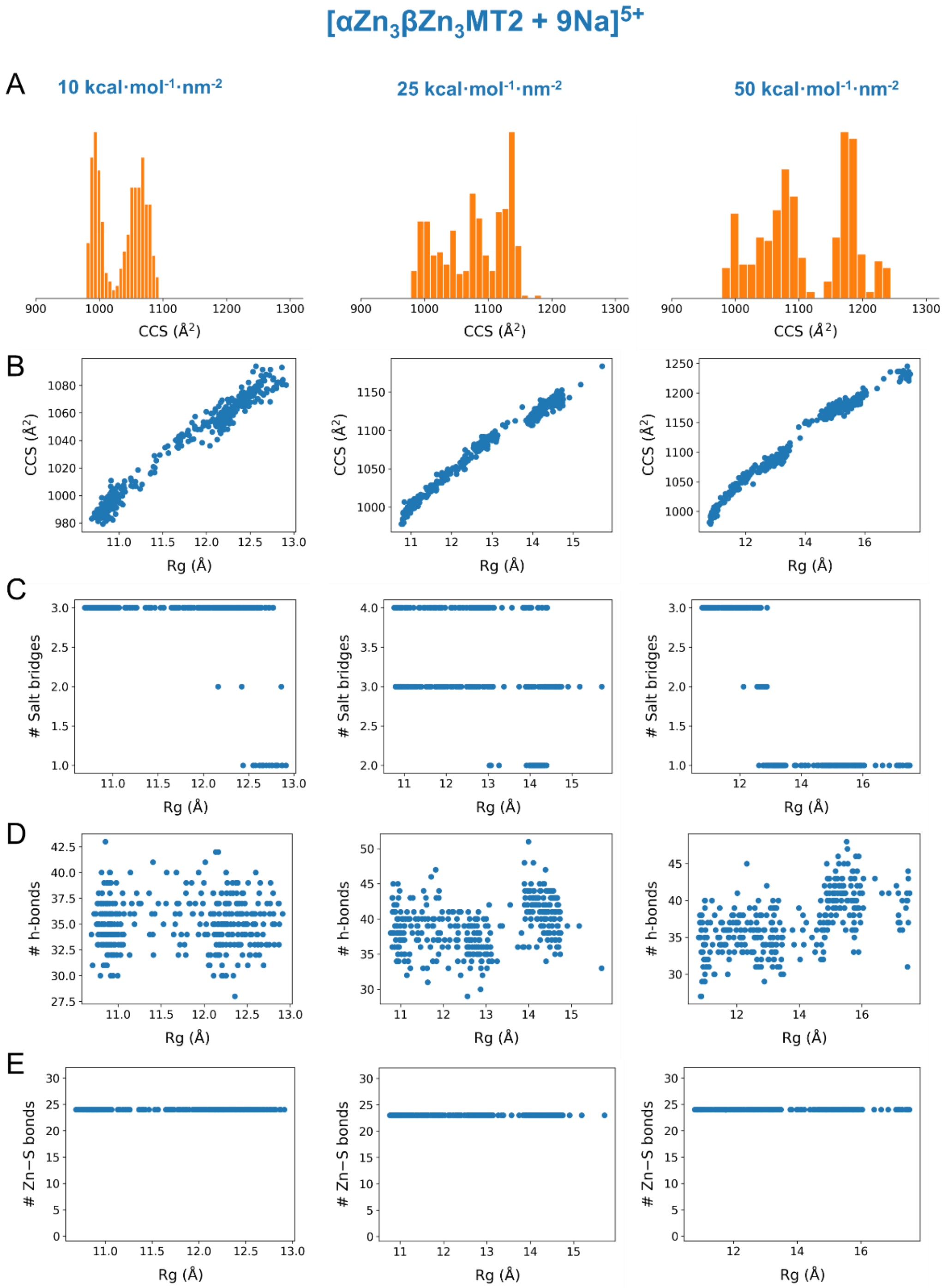


**Figure S15.** Optimization of the SMD simulations involving gas-phase [$\alpha Zn_3\beta Zn_3$MT2 +9Na]$^{5+}$ ions using radius of gyration ($R_g$) as a collective variable. CCS histograms for three different force constants (10, 25, and 50 kcal·mol$^{-1}$·nm$^{-2}$) (A), CCS (B), number of salt bridges (C), number of hydrogen bonds (D) and number of Zn−S bonds (E) as a function of the $R_g$.

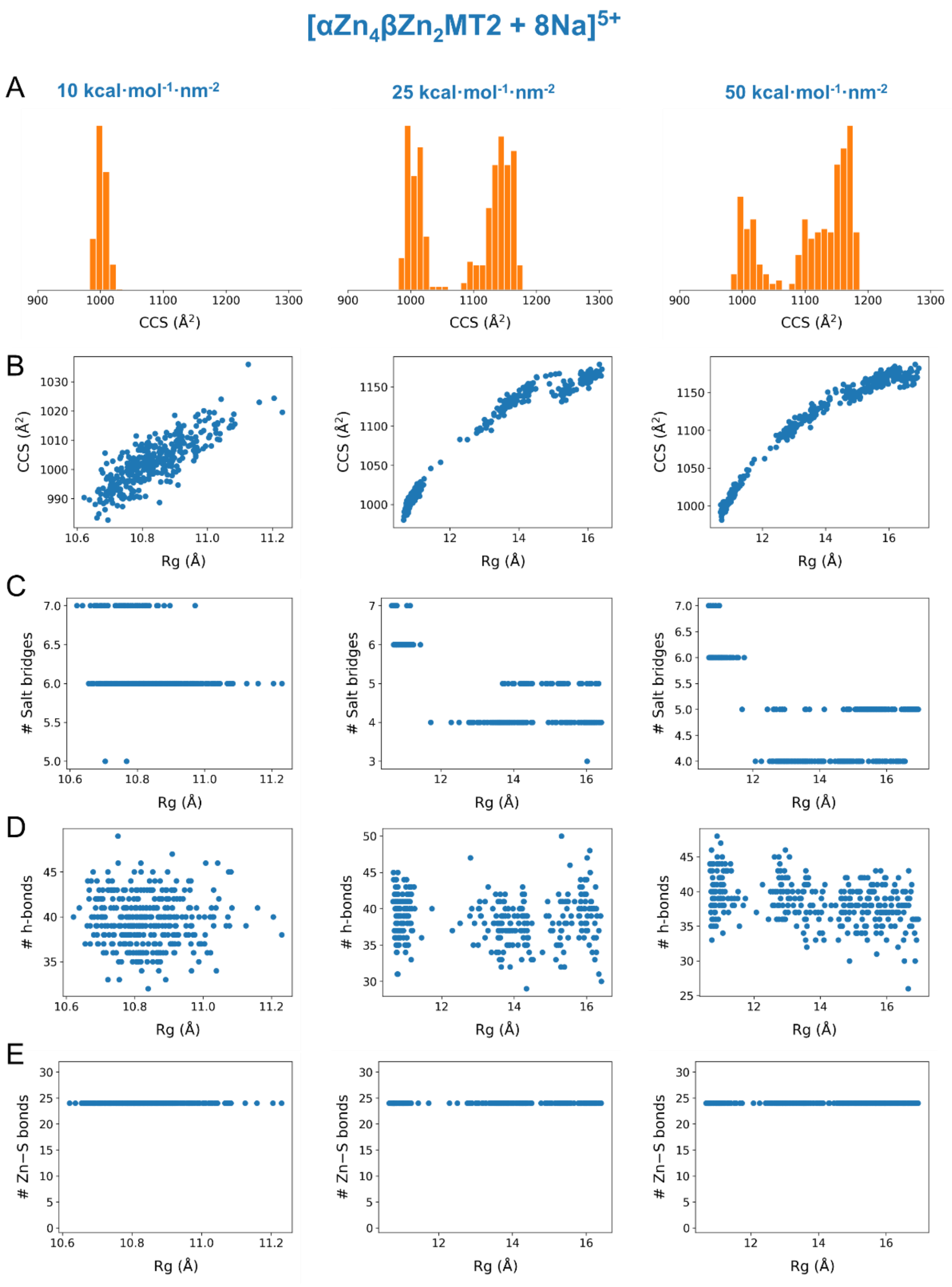


**Figure S16.** Optimization of the SMD simulations involving gas-phase [$\alpha Zn_4\beta Zn_2$MT2 +9Na]$^{5+}$ ions using radius of gyration ($R_g$) as a collective variable. CCS histograms for three different force constants (10, 25, and 50 kcal·mol$^{-1}$·nm$^{-2}$) (A), CCS (B), number of salt bridges (C), number of hydrogen bonds (D) and number of Zn−S bonds (E) as a function of the $R_g$.

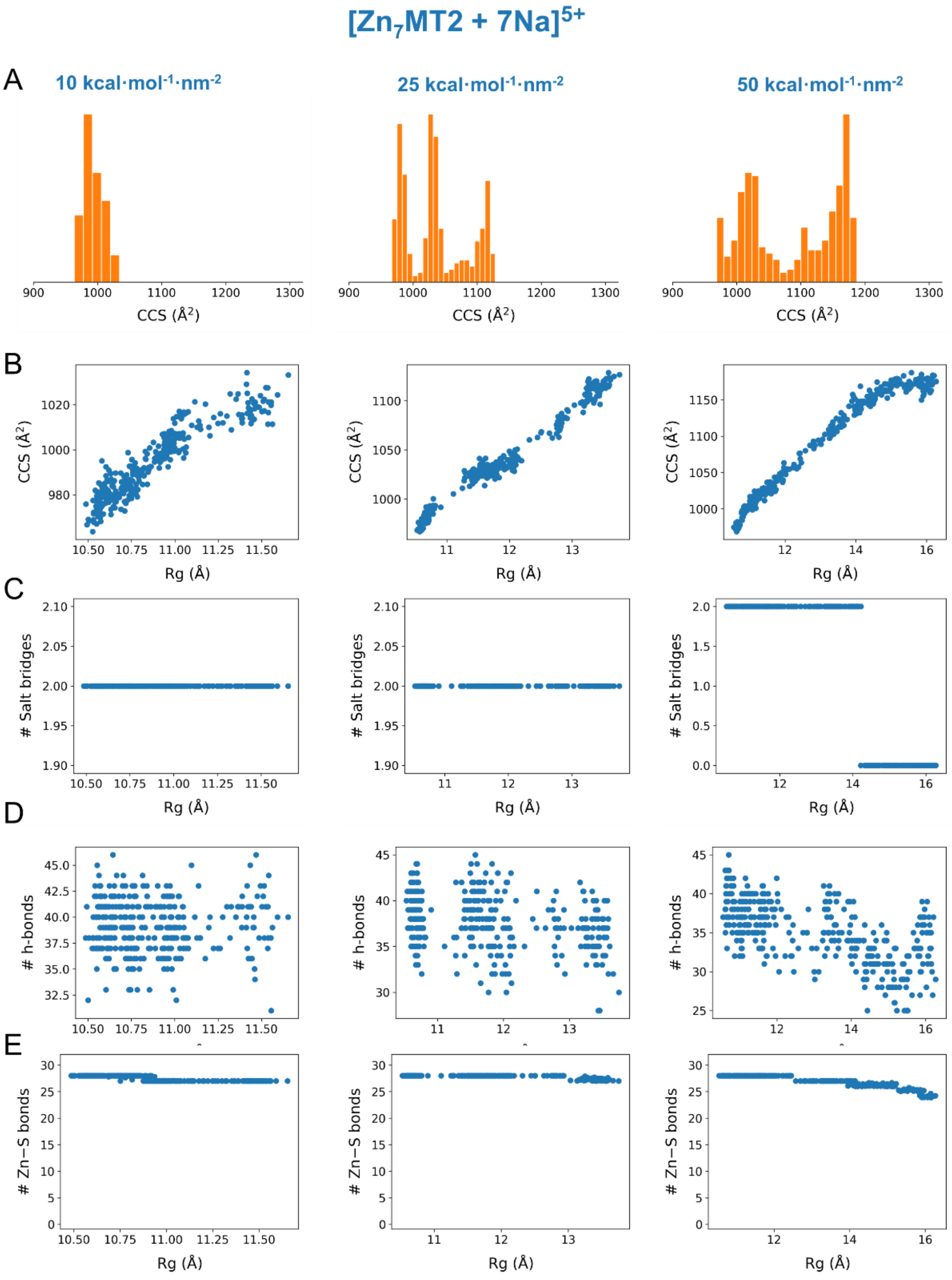


**Figure S17.** Optimization of the SMD simulations involving gas-phase [$Zn_7$MT2 +7Na]$^{5+}$ ions using radius of gyration ($R_g$) as a collective variable. CCS histograms for three different force constants (10, 25, and 50 kcal·mol$^{-1}$·nm$^{-2}$) (A), CCS (B), number of salt bridges (C), number of hydrogen bonds (D) and number of Zn−S bonds (E) as a function of the $R_g$.

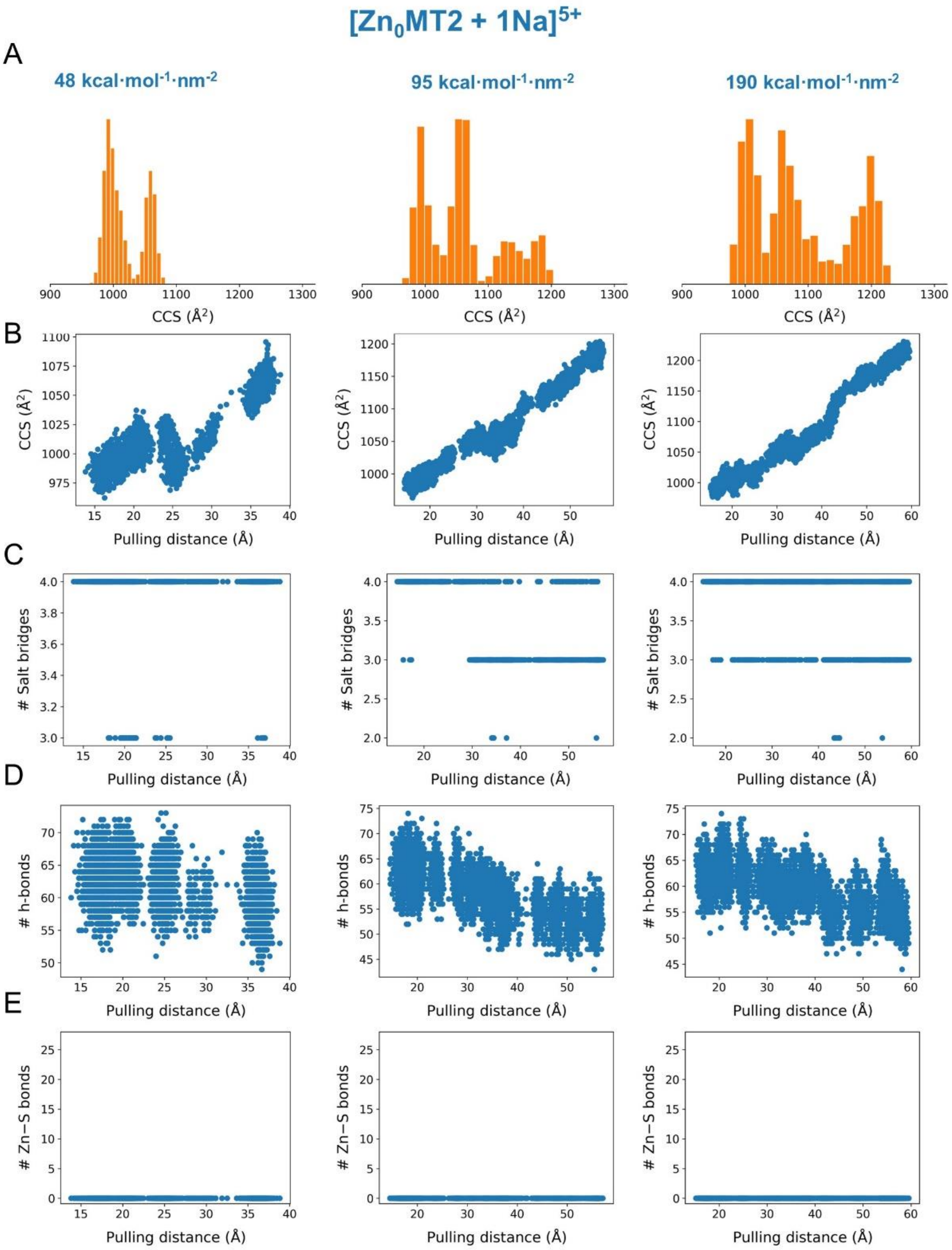


**Figure S18.** Optimization of the SMD simulations involving gas-phase [$Zn_0$MT2 + 1Na]$^{5+}$ ions using end-to-end distance (Pulling distance) as a collective variable. CCS histograms for three different force constants (48, 95, and 190 kcal·mol$^{-1}$·nm$^{-2}$) (A), CCS (B), number of salt bridges (C), number of hydrogen bonds (D) and number of Zn−S bonds (E) as a function of the Pulling distance.

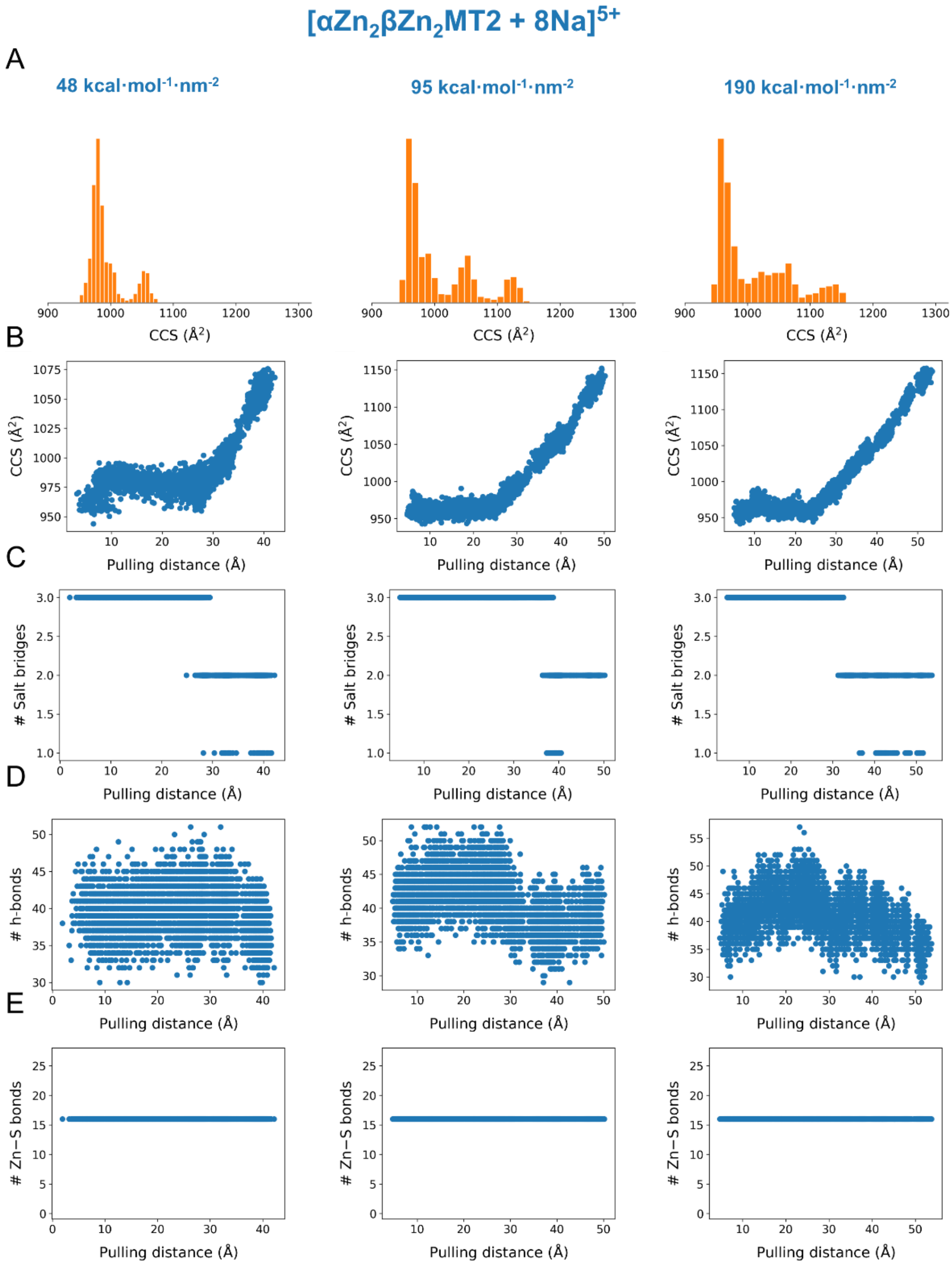


**Figure S19.** Optimization of the SMD simulations involving gas-phase [$\alpha Zn_2\beta Zn_2MT2$ +8Na]$^{5+}$ ions using end-to-end distance (Pulling distance) as a collective variable. CCS histograms for three different force constants (48, 95, and 190 kcal·mol$^{-1}$·nm$^{-2}$) (A), CCS (B), number of salt bridges (C), number of hydrogen bonds (D) and number of Zn−S bonds (E) as a function of the Pulling distance.

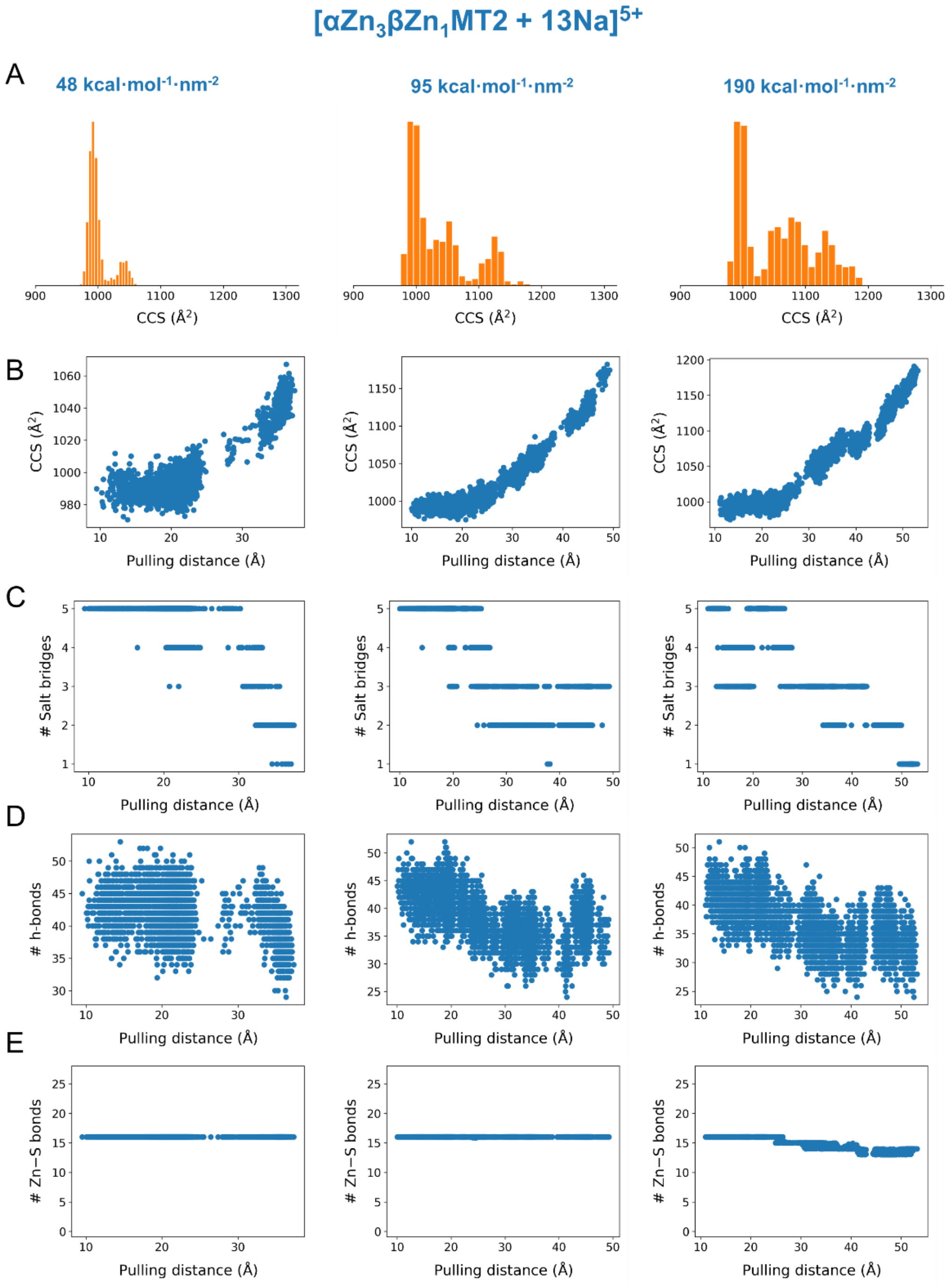


**Figure S20.** Optimization of the SMD simulations involving gas-phase [$\alpha Zn_3\beta Zn_1$MT2 +8Na]$^{5+}$ ions using end-to-end distance (Pulling distance) as a collective variable. CCS histograms for three different force constants (48, 95, and 190 kcal·mol$^{-1}$·nm$^{-2}$) (A), CCS (B), number of salt bridges (C), number of hydrogen bonds (D) and number of Zn−S bonds (E) as a function of the Pulling distance.

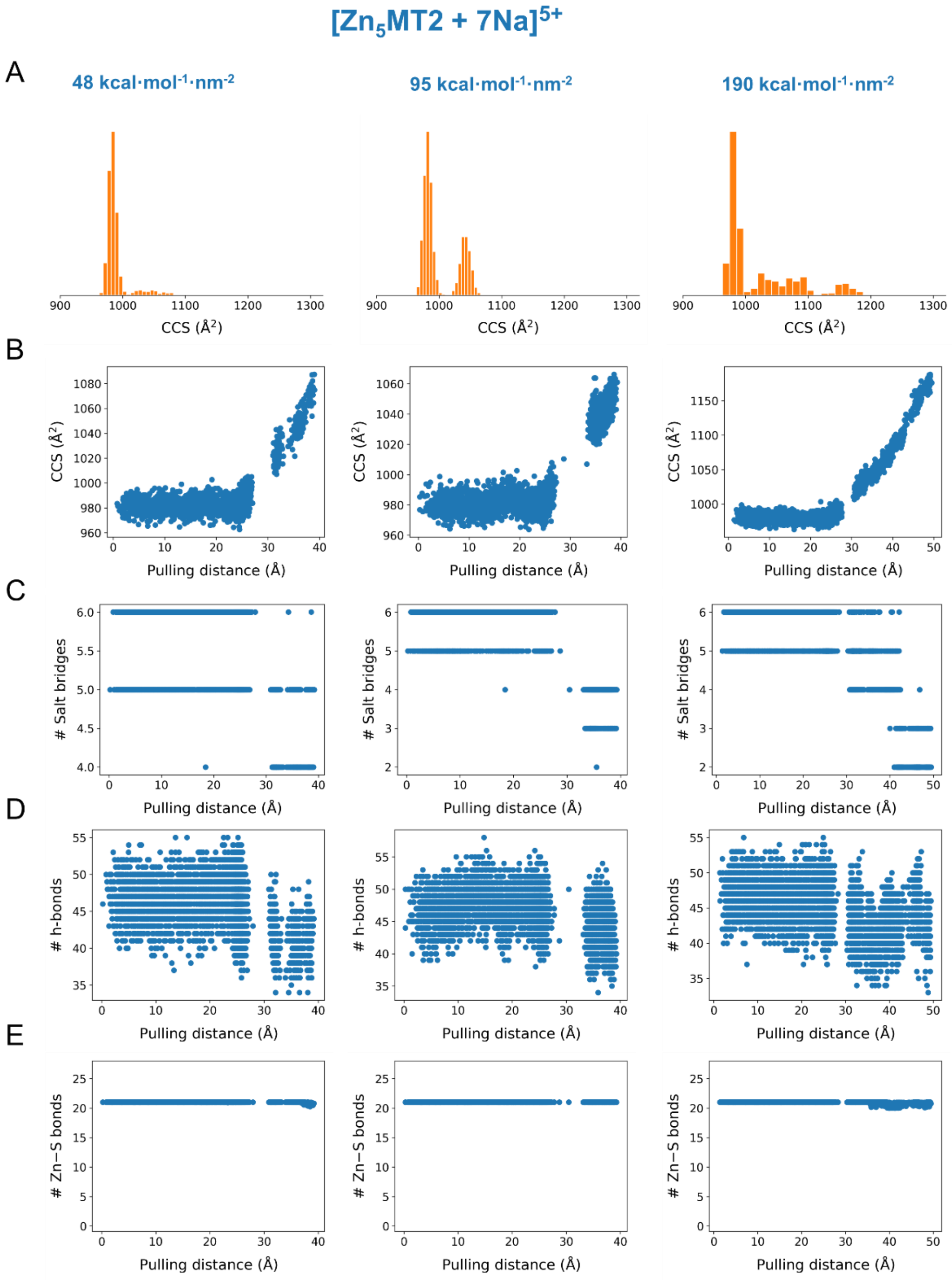


**Figure S21.** Optimization of the SMD simulations involving gas-phase $[Zn_5MT2 + 7Na]^{5+}$ ions using end-to-end distance (Pulling distance) as a collective variable. CCS histograms for three different force constants (48, 95, and 190 kcal·mol$^{-1}$·nm$^{-2}$) (A), CCS (B), number of salt bridges (C), number of hydrogen bonds (D) and number of Zn−S bonds (E) as a function of the Pulling distance.

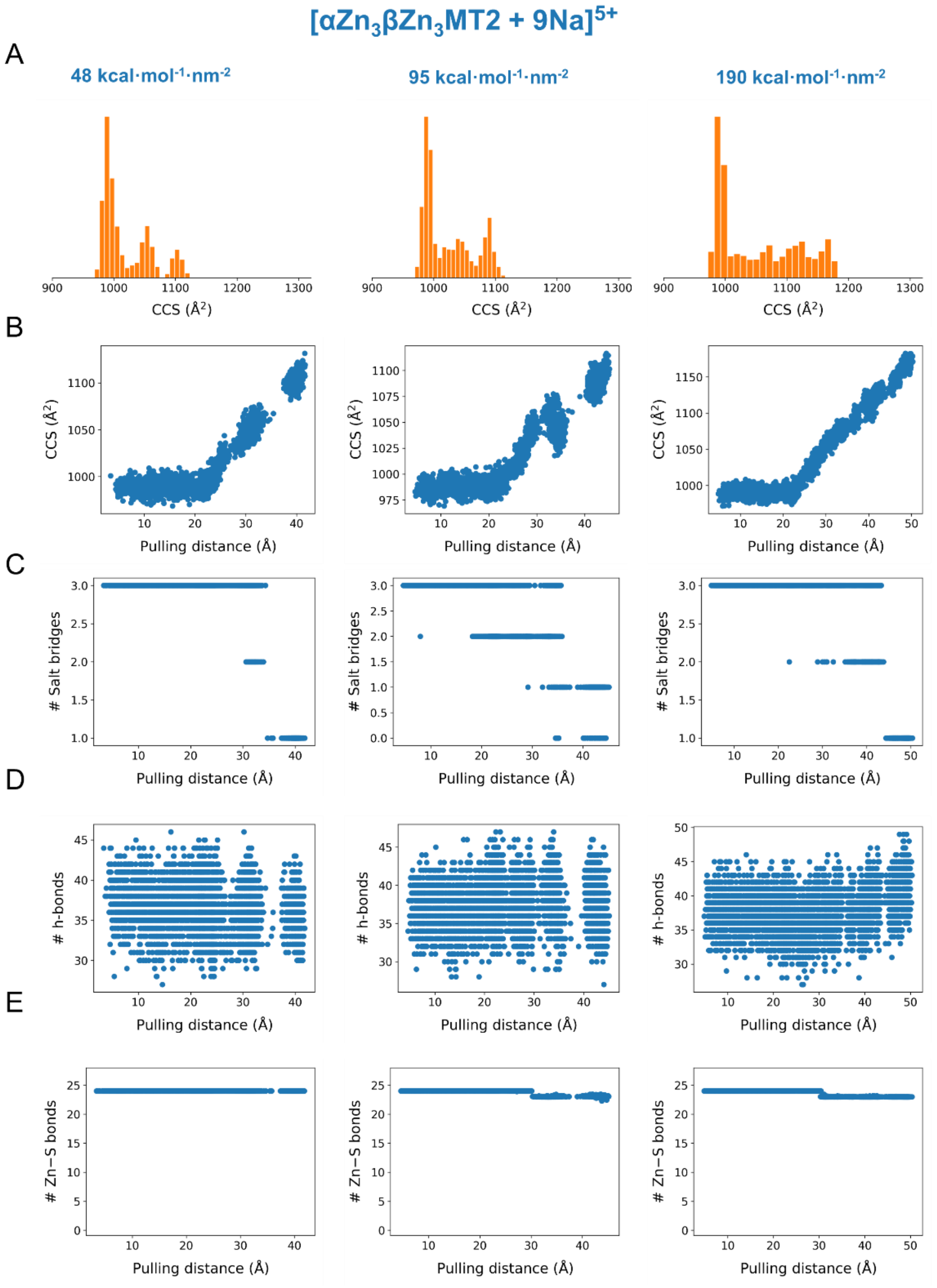


**Figure S22.** Optimization of the SMD simulations involving gas-phase $[\alpha Zn_3\beta Zn_3MT2 + 9Na]^{5+}$ ions using end-to-end distance (Pulling distance) as a collective variable. CCS histograms for three different force constants (48, 95, and 190 kcal·mol$^{-1}$·nm$^{-2}$) (A), CCS (B), number of salt bridges (C), number of hydrogen bonds (D) and number of Zn−S bonds (E) as a function of the Pulling distance.

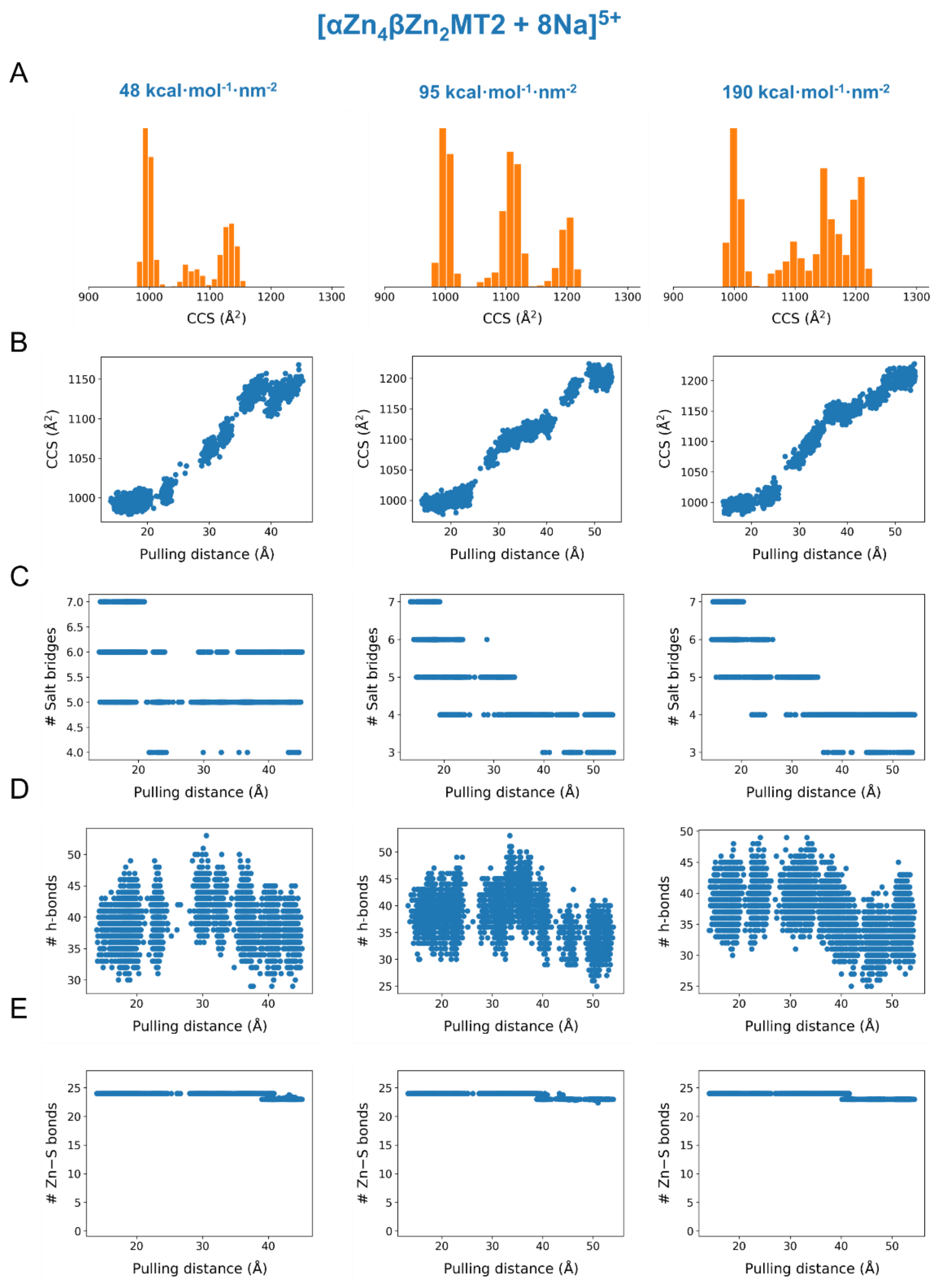


**Figure S23.** Optimization of the SMD simulations involving gas-phase [$\alpha Zn_4\beta Zn_2$MT2 + 8Na]$^{5+}$ ions using end-to-end distance (Pulling distance) as a collective variable. CCS histograms for three different force constants (48, 95, and 190 kcal·mol$^{-1}$·nm$^{-2}$) (A), CCS (B), number of salt bridges (C), number of hydrogen bonds (D) and number of Zn−S bonds (E) as a function of the Pulling distance.

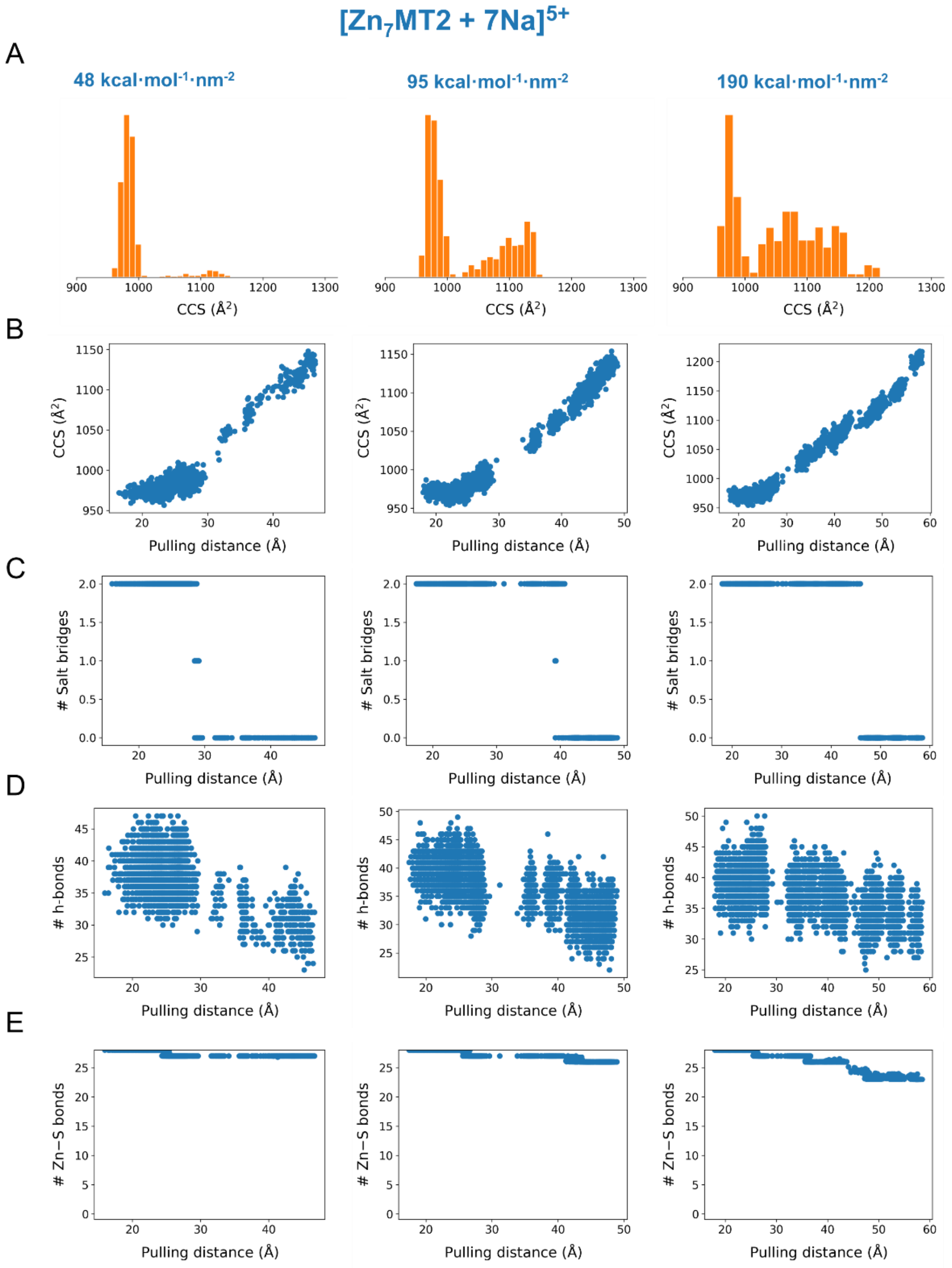


**Figure S24.** Optimization of the SMD simulations involving gas-phase $[Zn_7MT2 + 7Na]^{5+}$ ions using end-to-end distance (Pulling distance) as a collective variable. CCS histograms for three different force constants (48, 95, and 190 kcal·mol$^{-1}$·nm$^{-2}$) (A), CCS (B), number of salt bridges (C), number of hydrogen bonds (D) and number of Zn−S bonds (E) as a function of the Pulling distance.

**Table S1**. Accurate mass measurements of apoMT2 and $Zn_{4\text{-}7}MT2$ protein by native MS. The mass error corresponds to the difference between the experimental and the fitted molecular formula.

| Protein | Oxidation status | formula | Mass error (Da) | $\Delta H^{+}{}_{(apo\text{-}Zn\mathit{x}MT)\mathit{red}}$[a] | $\Delta H^{+}{}_{(apo\text{-}Zn\mathit{x}MT)\mathit{ox}}$[b] | $\Delta H^{+}{}_{red\text{-}ox}$[c] |
|---|---|---|---|---|---|---|
| apoMT2 | reduced | $C_{223}H_{381}O_{82}N_{71}S_{21}$ | 0.44 | 0 | - | - |
| | oxidized | $C_{223}H_{366}O_{82}N_{71}S_{21}$ | 0.12 | - | 0 | 15 |
| $Zn_4MT2$ | reduced | $C_{223}H_{371}O_{82}N_{71}S_{21}Zn_4$ | 0.24 | 10 | - | - |
| | oxidized | $C_{223}H_{354}O_{82}N_{71}S_{21}Zn_4$ | 0.32 | - | 12 | 17 |
| $Zn_5MT2$ | reduced | $C_{223}H_{365}O_{82}N_{71}S_{21}Zn_5$ | 0.02 | 16 | - | - |
| | oxidized | $C_{223}H_{349}O_{82}N_{71}S_{21}Zn_5$ | 0.25 | - | 17 | 16 |
| $Zn_6MT2$ | reduced | $C_{223}H_{365}O_{82}N_{71}S_{21}Zn_6$ | 0.22 | 16 | - | - |
| | oxidized | $C_{223}H_{349}O_{82}N_{71}S_{21}Zn_6$ | 0.06 | - | 17 | 16 |
| $Zn_7MT2$ | reduced | $C_{223}H_{367}O_{82}N_{71}S_{21}Zn_7$ | 0.25 | 14 | - | - |
| | oxidized | $C_{223}H_{359}O_{82}N_{71}S_{21}Zn_7$ | 0.01 | - | 7 | 8 |

[a]Stands for the number of protons dissociated of the reduced $Zn_xMT$ (x = 4-7) complex with respect to reduced apoMT2. [b]Stands for the number of protons dissociated from the oxidized $Zn_xMT$ (x = 4-7) complex with respect to oxidized apoMT2. [c]Stands for the number of protons dissociated of the reduced with respect to the oxidized $Zn_xMT$ (x = 4-7) complex.

**Table S2.** Summary of the protein systems and computational methods employed in this work.

| Systems | Method | Charge scheme | No. runs[a] | Simulation time (ns) |
|---|---|---|---|---|
| $Zn_{0,4\text{-}7}MT2$ | cMD | mobile $Na^+$ | 3 | 4200 |
| | | mobile $Na^+$/static $H^+$ | 3 | 4200 |
| $Zn_7MT2$ | | mobile $H^+$ | 2 | 3000 |
| $Zn_{0,4\text{-}7}MT2$ | desolvation | mobile $Na^+$ | 2 | 945 |
| $Zn_{0,4\text{-}7}MT2$ | SA | mobile $Na^+$ | 3 | 210 |
| | SMD (CV: $R_g$) | mobile $Na^+$ | 25 | 120 |
| | SMD (CV: N–C dis) | mobile $Na^+$ | 25 | 105 |
| | | | Total | 12780 |

[a] Stands for the number of independent runs of each protein system. Note that $Zn_{0,4\text{-}7}MT2$ refers to five protein systems that were simulated separately. Abbreviations: cMD, classical molecular dynamics; SA, simulated annealing; SMD, steered molecular dynamics; CV, collective variable; $R_g$, radious of gyration.
